# Epidermal Collagen Reduction Drives Selective Aspects of Aging in Sensory Neurons

**DOI:** 10.1101/2024.05.07.593041

**Authors:** Meera M. Krishna, Swapnil G. Waghmare, Ariel L. Franitza, Emily C. Maccoux, E Lezi

## Abstract

Despite advances in understanding molecular and cellular changes in the aging nervous system, the upstream drivers of these changes remain poorly defined. Here, we investigate the roles of non-neural tissues in neuronal aging, using the cutaneous PVD polymodal sensory neuron in *Caenorhabditis elegans* as a model. We demonstrate that during normal aging, PVD neurons progressively develop excessive dendritic branching, functionally correlated with age-related proprioceptive deficits. Our study reveals that decreased collagen expression, a common age-related phenomenon across species, triggers this process. Specifically, loss-of-function in *dpy-5* or *col-120*, genes encoding cuticular collagens secreted to the epidermal apical surface, induces early-onset excessive dendritic branching and proprioceptive deficits. Adulthood-specific overexpression of *dpy-5* or *col-120* mitigates excessive branching in aged animals without extending lifespan, highlighting the specific role of these collagens in promoting neuronal healthspan. Notably, collagen reduction specifically drives excessive branching in select sensory neuron subclasses but does not contribute to dendritic beading, another aging-associated neurodegenerative phenotype distinctively associated with a different mechanosensitive dysfunction. Lastly, we identify that *rig-3*, an Immunoglobulin Superfamily member expressed in interneurons, acts upstream of collagen genes to maintain PVD dendritic homeostasis during aging, with downstream requirement of *daf-16*/FOXO. These findings reveal that age-related collagen reduction cues neuronal aging independently of collagen’s traditional structural support function, potentially involving bi-directional communication processes between neurons and non-neuronal cells. Our study also offers new insights into understanding selective neuron vulnerability in aging, emphasizing the importance of multi-tissue strategies to address the complexities of neuronal aging.

## 1. INTRODUCTION

Age represents a significant risk factor for many neurological disorders, including peripheral neuropathies. While several factors such as diabetes, autoimmune diseases, physical injury, and vitamin imbalances contribute to neuropathies, aging not only increases susceptibility to these conditions but also raises the prevalence of idiopathic neuropathies (Hicks et al., 2021). Currently, our limited ability to treat neuropathies in elderly populations largely stems from gaps in understanding the mechanisms driving the aging of the nervous system. Signaling from non-neural tissues may be a promising avenue to address this gap.

Non-neural tissues, such as the skin, play essential roles in the development of the peripheral nervous system. Studies show that neurotrophic factors in the skin guide sensory innervation, with overexpression of neurotrophin-4 in mouse epidermal basal cells leading to excessive sensory endings in the dermal footpad (Krimm et al., 2006). Similarly in *C. elegans*, the morphogenesis of the PVD neuron, a cutaneous sensory neuron with an elaborate dendritic arbor, relies on SAX-7, an epidermally-expressed cell-adhesion molecule, along with other extracellular proteins, that precisely guide dendritic arborization (Dong et al., 2013; Salzberg et al., 2013). Epidermal ensheathment is a process where epidermal cells wrap around growing neurites of sensory neurons to regulate branching morphogenesis, which is also highly dependent on two-way signaling between the neuron and the skin; blocking sheath formation in *Drosophila* and zebrafish destabilizes dendrite branches and reduces nociceptive function (Jiang et al., 2019). However, whether non-neural tissues also actively contribute to the aging of the peripheral nervous system remains less understood.

As neurons age, they undergo structural deterioration, leading to synapse loss and impaired, signal transmission, and ultimately behavioral abnormalities. Neuritic beading, also known as focal swelling or blebbing, is an evolutionarily conserved aging-associated structural change indicative of neurodegeneration (Takeuchi et al., 2005). These neuritic beads, often consisting of broken-down cytoskeletal and motor protein components, disrupt organelle transport, such as mitochondrial trafficking, leading to bioenergetic dysfunction and eventual neuronal death (Takeuchi et al., 2005). Our previous research in *C. elegans* has shown that age-related upregulation of antimicrobial peptides (AMPs) in the skin triggers dendritic beading in PVD neurons, accompanied by mechanosensory decline, demonstrating a significant role for skin-derived molecular cues in sensory neuron aging (E et al., 2018).

Increased aberrant neurite branching is another age-related structural change, documented in the peripheral nervous systems of elderly human patients with sensory neuropathies (Lauria et al., 1999). While it has been speculated that the increased branching may compensate for other aging-associated structural and functional losses, this counterintuitive phenomenon is not well understood. Interestingly, *C. elegans* PVD neurons also exhibit a pronounced increase in dendritic branching in advanced age, as noted in our prior work (E et al., 2018) and in a separate study by Kravtsov et al. (Kravtsov et al., 2017), though the functional implications and precise underlying mechanisms remain unknown. It is noteworthy that the aging-associated excessive branching, unlike the dendritic beading, does not appear to be regulated by skin AMPs, suggesting the involvement of alternative mechanisms.

In this study, we provide an in-depth characterization of aging-associated excessive dendritic branching in PVD neurons, demonstrating its functional relevance to age-related decline in proprioception. Using PVD neuron as a model, we further discover that mutations or downregulation of two cuticular collagen genes, *dpy-5* and *col-120*, induce early-onset excessive dendritic branching and proprioceptive deficits, suggesting that age-dependent reduction of skin collagens plays a causative role in sensory neuron aging. Moreover, overexpression of these collagen genes mitigates the severity of excessive branching in aged animals. Importantly, our results also indicate that the regulatory role of skin collagens is specific to branching integrity during aging and is limited to certain types of cutaneous sensory neurons. Additionally, we identify RIG-3, a neuronal Immunoglobulin Superfamily protein, as an upstream partner in skin collagen’s regulation of PVD dendritic integrity, with downstream involvement of DAF-16/FOXO. These findings reveal a novel role for collagens in actively regulating sensory neuron aging, expanding our understanding of the upstream molecular mechanisms that drive nervous system aging.

## 2. RESULTS

### 2.1 Aging triggers progressive development of excessive higher-order dendritic branching in PVD sensory neurons

To investigate how skin tissues impact sensory neuron aging, we used the PVD neuron in *C. elegans* as a model. PVD is a cutaneous mechanosensory neuron, involved in harsh touch, cold sensation, and proprioception, extending dendrites across the body (Chatzigeorgiou et al., 2010; Tao et al., 2019). Each animal possesses two PVD neurons, PVDR on the right and PVDL on the left, with dendritic branches extending dorsally and ventrally. Each PVD has a single axon that synapses ventrally with AVA and PVC interneurons (Figure 1a) (Smith et al., 2010; White et al., 1986). From the PVD soma, two primary (1°) dendritic branches extend anteriorly and posteriorly. Secondary (2°) dendrites emerge orthogonally from 1° branches at regular intervals, forming non-overlapping ‘menorah-like’ repeating units mainly composed of tertiary (3°) and quaternary (4°) dendrites (Figure 1b) (Smith et al., 2010). Our previous studies have shown that aging in PVD neurons often involves the progressive development of bead-like structures along the dendrites (Figure S1a) (E et al., 2018), resembling morphological changes observed in human patients with peripheral neuropathies (Faerman et al., 1982). Majority of the beading structures in PVD dendrites indicate a disrupted microtubule network and are associated with age-related deficits in the harsh touch sensing function of PVD neurons (Figure S1b) (E et al., 2018).

**Figure 1:**
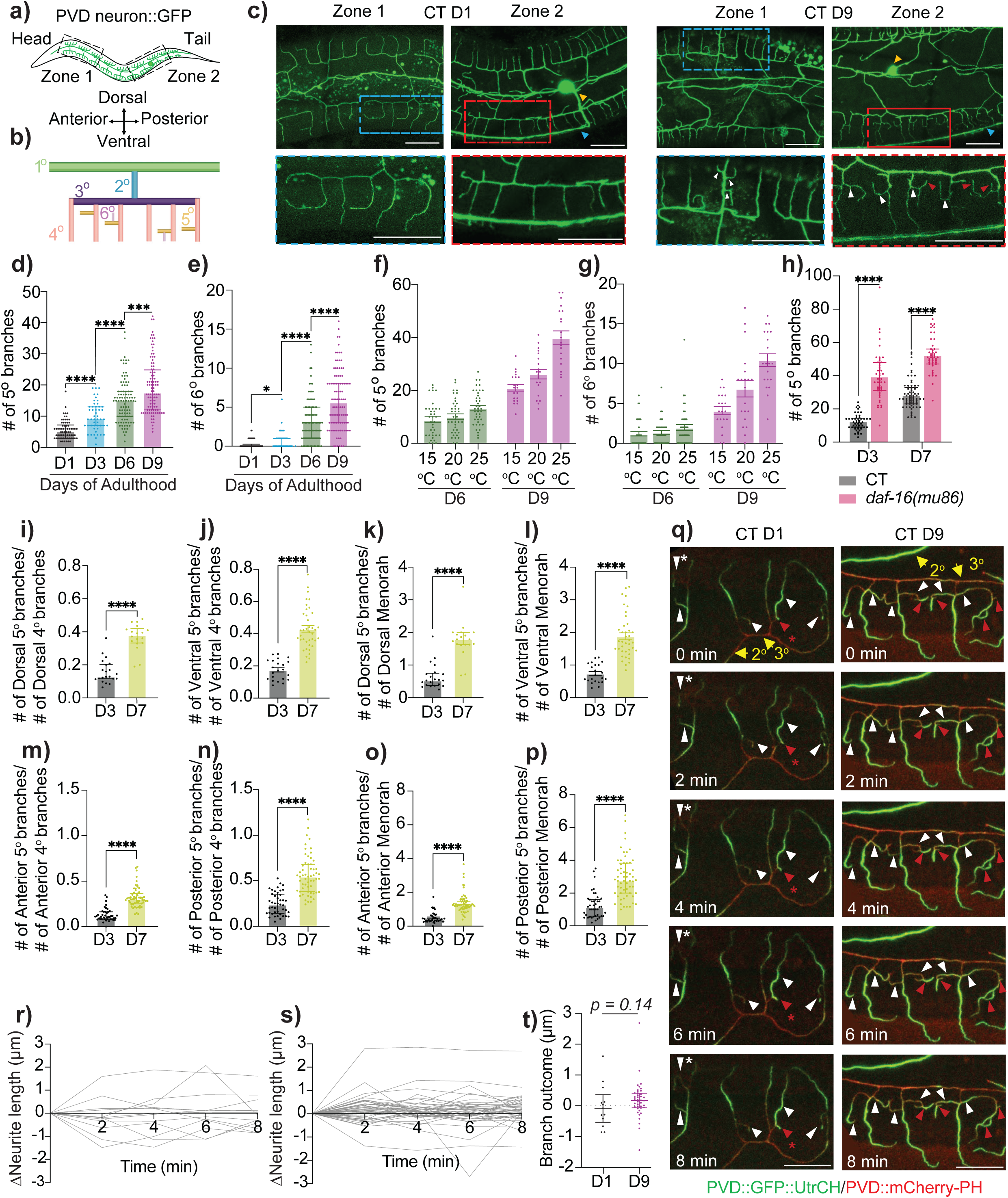
Aging triggers the progressive development of excessive higher-order dendritic branching in PVD neurons. **(a)** Schematic diagram of PVD neuron in control (CT) animals [*F49H12.4::GFP(wdIs51)*]. Black boxes indicate regions imaged. **(b)** A ‘menorah’ structure with orders of dendritic branching indicated. **(c)** Representative images of anterior (Zone 1) and posterior (Zone 2) sections of PVD neuron at Day 1 (D1) and D9 of adulthood in CT animals. Blue and red boxes indicate zoomed in sections for Zone 1 and 2 respectively. Yellow and blue arrowheads indicate PVD soma and axon respectively. White and red arrowheads indicate 5° and 6° dendritic branches respectively. Scale bar = 20 μM. Mild autofluorescence of gut granules, a normal physiological feature in *C. elegans*, appears as small punctate structures in the background. **(d-e)** Quantification of 5° *(d)* and 6° *(e)* dendrites during aging in CT animals (D1: n = 70, D3: n = 50, D6: n = 98, D9: n = 100). **(f-g)** 5° *(f)* and 6° *(g)* dendrites in CT animals cultured at different temperatures (D6 15° C: n = 36, D6 20° C: n = 39, D6 25° C: n = 41, D9 15° C, 20° C, 25° C: n = 20). **(h)** 5° dendrites in CT and *daf-16(mu86)* mutant animals (CT D3: n = 51, CT D7: n = 70, *daf-16* D3: n = 47, *daf-16* D7: n = 46). Experiments performed without FUDR. **(i-p)** Number of 5° dendrites normalized to number of 4° dendrites or menorahs. PVD soma was used as a reference point to determine anterior and posterior sections. *(i)* D3: n = 21, D7: n = 19. *(j)* D3: n = 22, D7: n = 39. *(k)* D3: n = 21, D7: n = 19. *(l)* D3: n = 22, D7: n = 39. *(m-p)* D3: n = 43, D7: n = 58. Experiments performed without FUDR. **(q)** Representative time-lapse images of actin dynamics in higher-order dendritic branches during an 8-minute session at 2-minute intervals in D1 and D9 CT animals [*ser-2(3)p::GFP::UtrCH + ser-2(3)p::mCherry-PH(lxyEx51)*]. mCherry::PH was used to visualize the membrane structures of PVD dendrites. Yellow arrows indicate 2° and 3° dendrites. White and red arrowheads indicate 5° and 6° dendrites, with asterisks indicating dendrites where a dynamic actin remodeling event occurs. Scale bar = 10 μm. **(r-s)** Changes in the length of higher-order dendrites relative to their initial lengths were tracked using time-lapse imaging data obtained from D1*(r)* and D9*(s)* CT animals as in *(q).* Measurements were taken for 5° and 6° dendrites branching from 4° and 5° dendrites, respectively, with each line representing a single higher-order dendrite. *n* (animals imaged) = 3/group. **(t)** Absolute change in branch length over 8 minutes in D1 and D9 CT animals (measurements from *(r-s)*), with each point representing a higher-order dendrite. n = 3/group. * *p*<0.05, *** *p*<0.001, **** *p*<0.0001.

In this study, we focused on and conducted a thorough characterization of another distinctive aging phenotype of PVD neurons – excessive dendritic branching, previously noted in literature (E et al., 2018; Kravtsov et al., 2017) yet lacking clarification regarding its functional implications and underlying mechanisms. Our analysis revealed a progressive and substantial increase in the number of 5° and 6° dendritic branches from Day 1 (D1) to D9 of adulthood in control (CT) animals (Figure 1c-e, Figure S1c-f), while the number of 4° dendrites or PVD menorahs remained mostly unchanged (Table S1). This phenotype was observed consistently regardless of FUDR use for synchronizing age (see Methods) (Table S1). Notably, no age-related ectopic branching was detected in the axons. To confirm that this branching is an inherent consequence of aging, we first utilized the environment temperature-lifespan correlation in *C. elegans*, where higher temperatures accelerate aging, while lower temperatures decelerate it (Klass, 1977). Our findings revealed a temperature-dependent variation in the severity and progression of PVD dendritic branching (Figure 1f-g). We further assessed branching in lifespan-affecting mutants, focusing on DAF-2/insulin-like signaling and mitochondrial pathway, given their well-established roles in aging (Ishii et al., 1998; Lee & Lee, 2022). In short-lived *daf-16(mu86)* mutants, excessive branching appeared earlier and progressed rapidly with age; *mev-1(kn1)* mutants with similarly short lifespans showed the same trend (Figure 1h, Figure S1g-i). Notably, long-lived *daf-2(e1370)* mutants displayed excessive branching similar to CT animals at D9, consistent with previous findings by Kravtsov et al. (Figure S1j).

Additionally, the increase in higher-order (≥ 5°) dendritic branching persisted after accounting for the slight age-related increase in body length (Figure S1m-n), indicating it is not merely growth-related. The extent and progression of branching were comparable between PVDL and PVDR throughout aging (Table S1). While slight variations were noted between the dorsal and ventral sides, the overall increase in branching remained consistent on both sides (Table S1). This pattern held even after normalizing for the number of PVD ‘menorah’ structures or 4° branches (Figure 1i-l, Table S1). Using the PVD soma position as a reference point, a similar increase was observed in both anterior and posterior sections (Figure 1m-p, Table S1), Together, these data suggest that excessive branching in PVD higher-order dendrites is an intrinsic feature of aging.

### 2.2 Cytoskeletal composition of higher-order branches in aging PVD neurons

During development, stabilized microtubules are often found in large primary dendrites, while actin is abundant in small higher-order dendrites or dendritic spines, supporting anatomical plasticity (Kaech et al., 2001). To examine the cytoskeletal composition of the aging-associated higher-order dendritic branches in PVD neurons, we used transgenic strains expressing *ser-2(3)p::EMTB::GFP* and *ser-2(3)p::GFP::UtrCH* to visualize polymerized microtubules and actin filaments, respectively, in PVD neurons (E et al., 2018). We found that microtubules were absent in >3° branches in both aged and young adult CT animals (Figure S2a). Time-lapse imaging analysis of young adult CTs revealed abundant F-actin in 4° dendrites and at the leading edges of occasional higher-order branches, with the excessive 5°/6° branches in aged CTs primarily consisting of F-actin as well (Figure 1q, Figure S2b). Further quantitative analysis indicated dynamic remodeling of F-actin-rich higher-order branches in both young and aged CTs, with frequent outgrowths and retractions (Figure 1r-s). Notably, during the same observation period, ≥ 5° branches in aged animals tended to gain length, while those in young adults appeared more tightly regulated from further growth (Figure 1t). This regulation is likely independent of pruning or phagocytosis, as no fluorescent neuronal debris was observed near remodeled regions. These findings suggest that a localized mechanism restricting dendritic growth may be lost with aging.

### 2.3 Aging-associated excessive dendritic branching in PVD neurons correlates with proprioceptive deficits

To explore the functional implications of the excessive branching in aging PVD neurons, we first examine its relevance to noxious harsh touch sensation, a process relying on proper PVD synaptic connections with downstream command interneurons for initiating escape movements (Chatzigeorgiou et al., 2010; E et al., 2018). To correlate neuronal morphological changes with functional outcomes, we performed harsh touch assays with individual animals and subsequently analyzed PVD branching in the same animals. Interestingly, age-related decline in harsh touch response in CTs (E et al., 2018) showed no correlation with excessive branching (Figure S3a).

As a polymodal neuron, PVD senses multiple stimuli. The integrity of its dendritic menorah structures is essential for the sensory aspect of proprioceptive regulation, independent of its axon’s synaptic function (Tao et al., 2019). Proprioception, the processing of sensorimotor input for posture control and movement, can be assessed in *C. elegans* by measuring track patterns left by the worms on bacterial lawns (Figure 2a). Mutations in *dma-1* or related genes disrupting PVD menorah structures (characterized by absence of ≥ 2° branches) reduce track wavelength and amplitude (Tao et al., 2019). We therefore hypothesized that excessive branching in aging PVD neurons could manifest as proprioceptive deficits.

**Figure 2:**
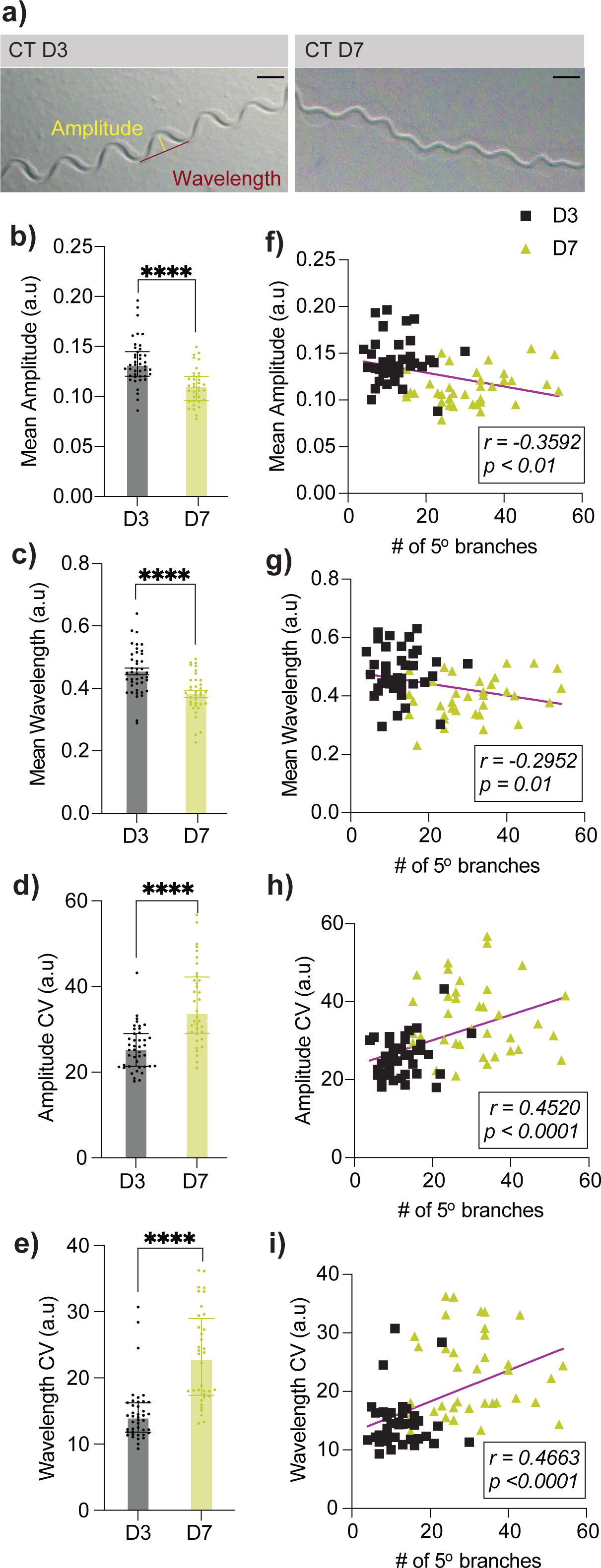
Aging-associated excessive dendritic branching in PVD neurons correlates with proprioceptive deficits. **(a)** Representative images of movement tracks of CT animals at D3 and D9. Scale bar = 500 μM. **(b-c)** Mean amplitude *(b)* and wavelength *(c)* measured from 100 tracks per animal in CT animals. Normalized for animal body length (D3: n = 42, D7: n = 36). **(d-e)** Variability in amplitude *(d)* and wavelength *(e)* measured via coefficient of variability (CV) from 100 tracks per animal in CT animals (D3: n = 42, D7: n = 36). **(f-i)** Correlation between proprioception measurements and number of PVD 5° dendritic branches. Each data point represents an individual animal. **** *p*<0.0001. All experiments in this figure performed without FUDR.

Our data showed that both the mean amplitude and mean wavelength of sinusoidal movements were significantly reduced in aged CT animals, compared to young (Figure 2b-c). Additionally, aged CT animals displayed increased intra-individual variability in both amplitude and wavelength (Figure 2d-e). These age-related changes resemble the gait pattern changes observed in elderly humans, such as decreased stride length/step width and greater variability, which are strongly linked to a decline in proprioception (Callisaya et al., 2010; Wiesmeier et al., 2015). A within-individual correlation analysis further showed that the extent of ‘higher-order’ branches, particularly dorsal branches, was negatively correlated with mean amplitude and wavelength, while the severity of excessive branching, regardless of orientation, positively correlated with variability in track amplitude and wavelength (Figure. 2f-i, Table S2). These findings reveal a novel link between aging-associated excessive branching of PVD dendrites and proprioceptive deficits in aging.

Interestingly, there appears to be an absence of concurrence of PVD dendritic beading and excessive branching within individual animals; rather, a negative correlation seems evident, with animals showing pronounced beading less likely to exhibit severe branching, and vice versa (Figure S3b-c). Combined with the distinct behavioral manifestations of beading and excessive branching (E et al., 2018) (Figure 2, Figure S1b, Figure S3a), these observations suggest that aging neurons undergo a variety of morphological changes, each linked to specific functional outcomes, potentially as a compensatory mechanism to preserve certain functions especially in polymodal neurons.

### 2.4 Loss-of-function mutations in epidermal collagen genes induce early-onset excessive PVD dendritic branching and proprioceptive deficits

We next investigated the molecular mechanisms underlying aging-associated excessive branching. Previously, we found that aging-associated upregulation of a skin AMP contributes to PVD dendritic beading during aging, indicating that age-related skin changes can influence PVD neuron aging (E et al., 2018). However, AMP upregulation does not account for the excessive branching phenotype (E et al., 2018), prompting us to explore whether other molecular changes in the aging skin might contribute. A prime candidate is age-related decline in skin collagen gene expression, a phenomenon well-documented across species, including humans. Among the downregulated collagen genes in aging *C. elegans* (Figure S4a, GSE176088) (Teuscher et al., 2024), we examined several, including *col-120*, *col-129*, *col-141*, *dpy-5*, *dpy-10* and *rol-6*, previously studied for roles in morphogenesis and other contexts. We found that disruption of *col-120*, *col-141*, *dpy-5* and *dpy-10* via RNA interference (RNAi) or mutation resulted in early-onset excessive branching in PVD dendrites by D3, while the others had no effects (Figure 3a-i; Figure S4b-i), suggesting specific skin collagens may play a role in neuronal aging. In this study, we focused on *col-120* and *dpy-5*, as neither impacts skin integrity or AMP gene expression, unlike *col-141* and *dpy-10* (Sandhu et al., 2021; Zhu et al., 2023), which could confound PVD aging analysis (E et al., 2018). Furthermore, the expression of *dpy-5* and *col-120* are not temporally clustered, and their proteins localize to distinct areas within the epidermal apical surface (Teuscher et al., 2024; Thacker et al., 2006), suggesting that they likely do not interact with each other functionally.

**Figure 3:**
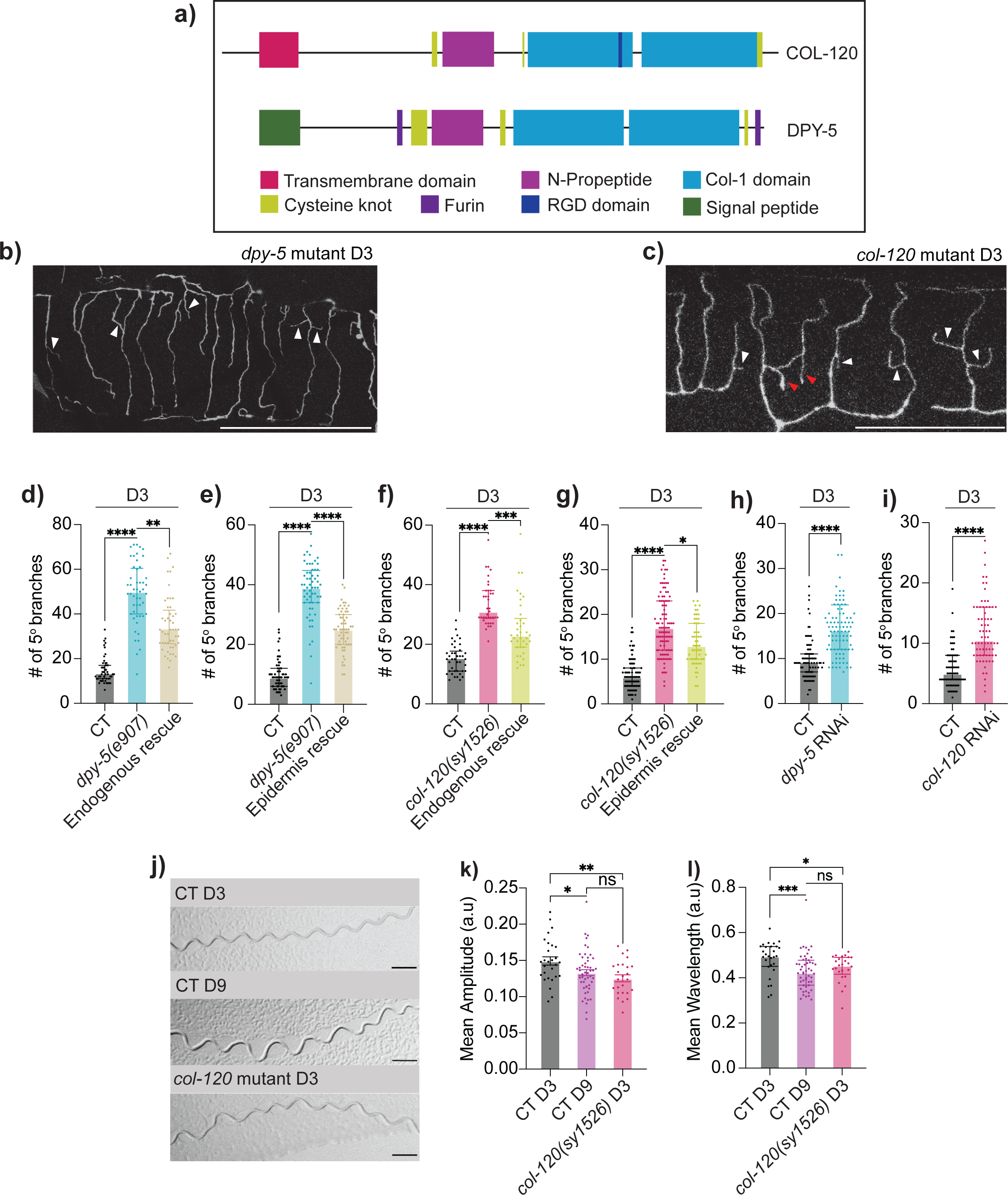
Loss of epidermal collagen leads to early-onset of aging-associated excessive branching and proprioceptive deficit. **(a)** Domain structures of COL-120 and DPY-5, with key features annotated. Adapted from CeCoIDB (Teuscher et al., 2019). **(b-c)** Representative images of excessive PVD dendritic branching in *dpy-5(e907)* mutant *(b)* and *col-120(sy1526)* mutant *(c)* animals at D3. White and red arrowheads indicate 5° and 6° dendrites respectively. Scale bar = 50 μM. **(d)** Quantification of 5° dendrites in CT (n = 50), *dpy-5* mutant (n = 50), and endogenous promoter-driven rescue in *dpy-5* mutant (n = 49). **(e)** 5° dendrites in CT, *dpy-5* mutant, and epidermis-specific rescue in *dpy-5* mutant (driven by *dpy-7p*; line 1) (n = 60 for all groups). **(f)** 5° dendrites in CT (n = 40), *col-120* mutant (n = 38), and endogenous promoter-driven rescue in *col-120* mutant (n = 38). **(g)** 5° dendrites in CT (n = 82), *col-120* mutant (n = 100), and epidermis-specific rescue in *col-120* mutant (driven by *dpy-7p* and *col-19p*; line 1) (n = 55). **(h)** 5° dendrites in empty vector (EV) (n = 70) and *dpy-5* RNAi treated animals (n = 88; RNAi clone 1). **(i)** 5° dendrites in EV (n = 70) and *col-120* RNAi treated animals (n = 70; RNAi clone 1). **(j)** Representative images of movement tracks of D3 CT, D9 CT and D3 *col-120(sy1526)* animals. Scale bar = 500 μM. **(k-l)** Mean amplitude *(k)* and wavelength *(l)* measured from 100 tracks per animal for D3 CT (n = 28), D9 CT (n = 51), and D3 *col-120* mutant (n = 25). Normalized for animal body length. ns-not significant, * *p*<0.05, ** *p*<0.005, *** *p*<0.001, **** *p*<0.0001.

In *C. elegans*, the skin is primarily composed of a simple epithelium that secretes a collagenous cuticle to the apical side as a surface coat, creating an additional protective barrier (Chisholm & Hsiao, 2012). DPY-5 and COL-120 are classified as cuticular collagens (Teuscher et al., 2019; Thacker et al., 2006), with a domain structure similar to mammalian membrane-associated or fibril-associated collagens with interrupted triple-helices, distinguishing them from basement membrane collagens (Figure 3a). Importantly, DPY-5 and COL-120 protein levels both decrease with age in wild-type animals (Teuscher et al., 2024; Thacker et al., 2006). Although mutations in either *dpy-5* or *col-120* do not affect skin integrity (Sandhu et al., 2021; Van de Walle et al., 2019), these mutants have reduced lifespans and altered body sizes, with *dpy-5(e907)* being shorter and wider and *col-120(sy1526)* slightly elongated (Figure S4k-m). To account for body size differences, we normalized PVD dendritic branching to individual body length, confirming that early-onset excessive branching persisted in both *col-120(sy1526)* and *dpy-5(e907)* at D3 (Figure S1k-n).

We further performed rescue experiments by expressing wild-type *dpy-5* or *col-120* cDNA under their respective endogenous promoters or an epidermis-specific promoter. Both approaches successfully rescued the excessive branching phenotype in each mutant (Figure 3d-g, additional rescue lines in Figure S5). PVD neurons complete dendritic arborization around the late L4 stage (last/4^th^ larval stage) (Smith et al., 2010). At L4, *dpy-5* and *col-120* mutants showed a slight increase in dendritic branching compared to CT animals, though not reaching the extent observed at D3 (Figure S5i-j). To confirm that excessive branching in *dpy-5* or *col-120* mutants is genuinely aging-associated, we performed adulthood-specific systemic RNAi knockdowns of these genes, initiating at late L4 and phenotyping PVD dendrites 3 days into adulthood (D3). With over 180 cuticular collagen genes in *C. elegans* (Wormbase), multiple predicted paralogs exist for both *col-120* and *dpy-5*. To ensure specificity, our RNAi clones targeted unique coding regions without homologous sequences in other collagen genes (see Methods). RNAi effectiveness was validated by RT-qPCR (Figure S4j). CT animals fed on HT115(DE3) bacteria with empty vector L4440 (EV) exhibited aging-associated excessive branching in PVD similar to those fed on OP50 (Figure S5k-l). Our adulthood-specific RNAi approach was sufficient to induce excessive higher-order branches by D3 (Figure 3h-i, Figure S5).

Importantly, we found that D3 *col-120(sy1526)* mutants exhibited a significant decrease in both mean wavelength and amplitude of sinusoidal movement, closely resembling CT animals at D9, though without increased variability in these parameters (Figure 3j-l, Figure S5t-u). Collectively, these findings demonstrate that epidermal collagens, DPY-5 and COL-120, play a critical role in maintaining PVD dendritic branching integrity and PVD-navigated proprioceptive function during aging.

### 2.5 Adulthood-specific overexpression of epidermal collagen genes delays onset of excessive PVD dendritic branching

Using the *col-19* promoter (Thein et al., 2003) we overexpressed *col-120* or *dpy-5* in the epidermis, specifically during adulthood, and found that it mitigated excessive branching in PVD when examined at D9 (Figure 4a-b, Figure S6a-d). Moreover, *dpy-5* overexpression appeared to shift proprioceptive function parameters in D9 animals towards those of younger animals, though this effect did not reach statistical significance (Figure S6f-i). It is important to note that age-related declines in other systems, such as muscles and motor neurons, may complicate assessing sensory aspect of proprioception at this stage of life. Interestingly, neither *dpy-5* nor *col-120* overexpression extended lifespan (Figure 4c-d, Figure S6e). Previous studies reported a slight lifespan extension with *col-120* overexpression (Ewald et al., 2015; Goyala & Ewald, 2023; Teuscher et al., 2024); however, this discrepancy may stem from differences in promoters and 3’UTR sequences used to construct transgenic strains. Nevertheless, our findings suggest that the effects of epidermal collagen overexpression on aging primarily target neuronal health, rather than through promoting longevity.

**Figure 4:**
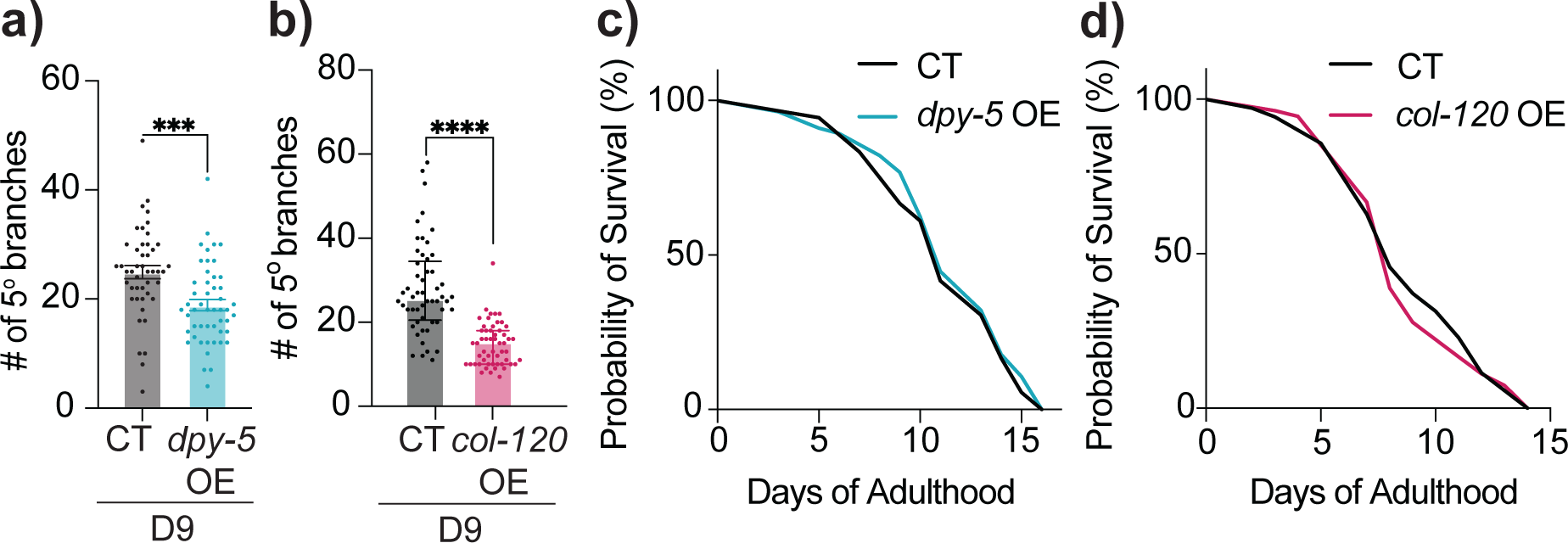
Adulthood-specific overexpression of collagen genes mitigates aging-associated excessive dendritic branching in PVD neurons. **(a)** Quantification of 5° dendrites in CT (n = 46) and *dpy-5* epidermis-specific overexpression driven by *col-19p* (n = 50). **(b)** 5° dendrites in CT (n = 53) and *col-120* epidermis-specific overexpression driven by *col-19p* (n = 54; line 2). **(c-d)** Lifespan comparison between CT (n = 100) and *dpy-5* epidermis-specific overexpression driven by *col-19p* (n = 100) *(c)*, as well as CT (n = 100) and *col-120* epidermis-specific overexpression driven by *col-19p* (n = 100; line 1) *(d)*. No statistical significance compared to CT. Additional replicates in Figure S6. Experiments performed without FUDR. *** *p*<0.001, **** *p*<0.0001.

### 2.6 DPY-5 and COL-120 are not involved in aging-associated PVD dendritic beading

To investigate whether skin collagen genes regulate the overall aging of PVD neurons, we examined the effects of *dpy-5* or *col-*120 mutation on the previously documented aging-associated dendritic beading phenotype (Figure S1a) (E et al., 2018). Notably, the extent of PVD dendritic beading in Day 3 *dpy-5(e907)* or *col-120(sy1526)* mutant animals was comparable to that observed in age-matched CT (Figure 5a). Coupled with our observations of a potential inverse relationship between aging-associated dendritic beading and excessive dendritic branching in older CT animals (Figure S3b-c), these results suggest that *dpy-5* and *col-120* regulate PVD dendritic structure via mechanisms separate from those leading to the dendritic beading degenerative phenotype, such as the skin AMP NLP-29 (E et al., 2018). This is also supported by previous research showing that mutations in *dpy-5* and *col-120* do not impact *nlp-29* expression (Zhu et al., 2023). Collectively, our findings support the notion that distinct non-neural molecular mechanisms may be underlying different dimensions of neuronal aging, even within an individual neuron.

**Figure 5:**
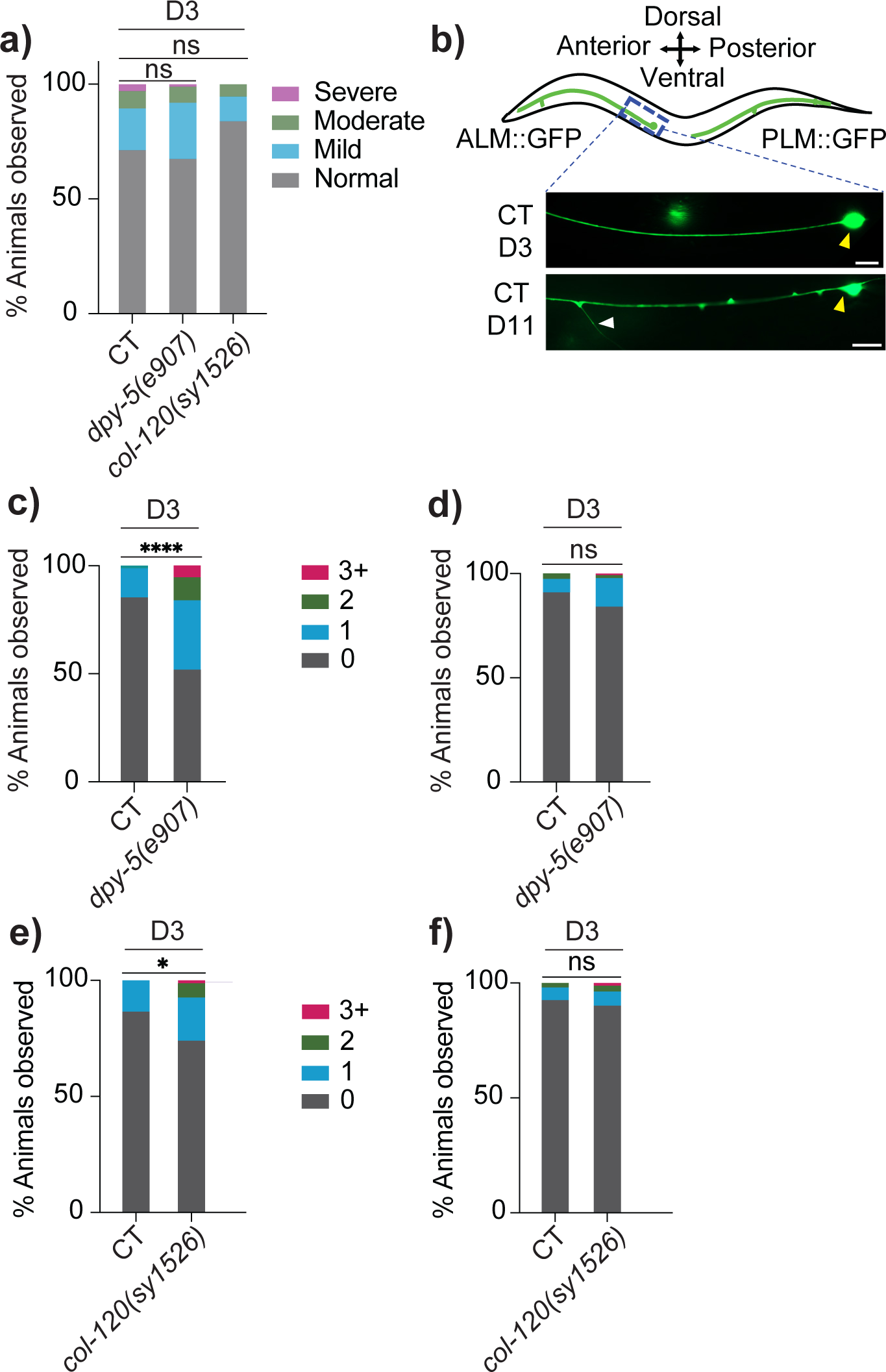
Selective impact of epidermal collagens on the aging process of sensory neurons. **(a)** Quantification of PVD dendritic beading in control (CT, n = 29), *dpy-5(e907)* (n = 26), and *col-120(sy1526)* (n = 29). Scored based on severity of beading phenotype. **(b)** Schematic diagram of ALM and PLM neuron, with a representative image of the ALM neuron in CT [*mec-4p::GFP(zdIs5)*] animals at D3 and D11. Scale bar = 50 μM. White and yellow arrowheads indicate the ectopic branching from ALM anterior neurite and the ALM soma respectively. **(c-d)** Ectopic branches in ALM *(c)* and PLM neuron *(d)* in CT (n = 78) and *dpy-5(e907)* (n = 131). Number of ectopic branches observed per animal is indicated. **(e-f)** Ectopic branches in ALM *(e)* and PLM neuron *(f)* in CT (n = 67) and *120(sy1526)* (n = 81). ns-not significant, * *p*<0.05, **** *p*<0.0001.

### 2.7 ALM sensory neurons exhibit ectopic branching in response to loss-of-function mutation in *dpy-5* or *col-120*, but PLM neurons do not

To determine whether the regulatory role of skin collagens in neuron structural integrity is specific to certain neuron populations, we examined two additional sub-types of cutaneous sensory neurons in *C. elegans*: ALM and PLM touch receptor neurons, which are crucial for the perception of gentle touch (Corsi et al., 2015). Intriguingly, both *dpy-5(e907)* and *col-120(sy1526)* animals exhibited an increase in ectopic, aberrant branching from ALM anterior neurites at D3 when compared to CT animals, while PLM neurons showed no morphological abnormalities (Figure 5b-f). This discrepancy indicates that the regulatory role of examined collagens in neuritic branching may not extend uniformly across all cutaneous sensory neurons, suggesting that certain neuron sub-types may possess unique features rendering them particularly vulnerable to the effects of skin collagen loss during aging.

### 2.8 RIG-3/IgSF in interneurons acts upstream of epidermal collagens to regulate excessive PVD dendritic branching in aging

Next, we explored the downstream mechanisms by which collagen genes regulate PVD dendritic integrity during aging. In *C. elegans*, the polarized structure of epidermal cells segregates apically secreted cuticular collagens from basement membrane collagens anatomically and functionally (Chisholm & Xu, 2012), making direct interaction between COL-120 or DPY-5 and sensory neurons (located on the basal side of the epidermis) unlikely. Thus, the age-related decline in these collagens may contribute to neuronal aging through indirect mechanisms, potentially involving cell signaling rather than direct structural support. To test this hypothesis, we first examined several neuronal cell surface receptors using adulthood-specific RNAi. We found that knockdown of *ina-1*, *pat-2* or *rig-3* led to early-onset excessive dendritic branching in PVD (Figure 6a, Figure S7a-c). While *ina-1* and *pat-2* (both α-integrins) are widely expressed, *rig-3*, encoding an Immunoglobulin Superfamily (IgSF) protein, is expressed selectively in a small set of neurons but notably absent in PVD (Babu et al., 2011; Taylor et al., 2021). This selective expression pattern makes *rig-3* a particularly intriguing candidate for a non-cell-autonomous role in regulating PVD morphology. Analysis of *rig-3(ok2156)* mutants confirmed our RNAi findings of early-onset excessive dendritic branching but showed that *rig-3* mutation does not significantly impact PVD morphogenesis during development (Figure 6b, Figure S7d-f).

**Figure 6:**
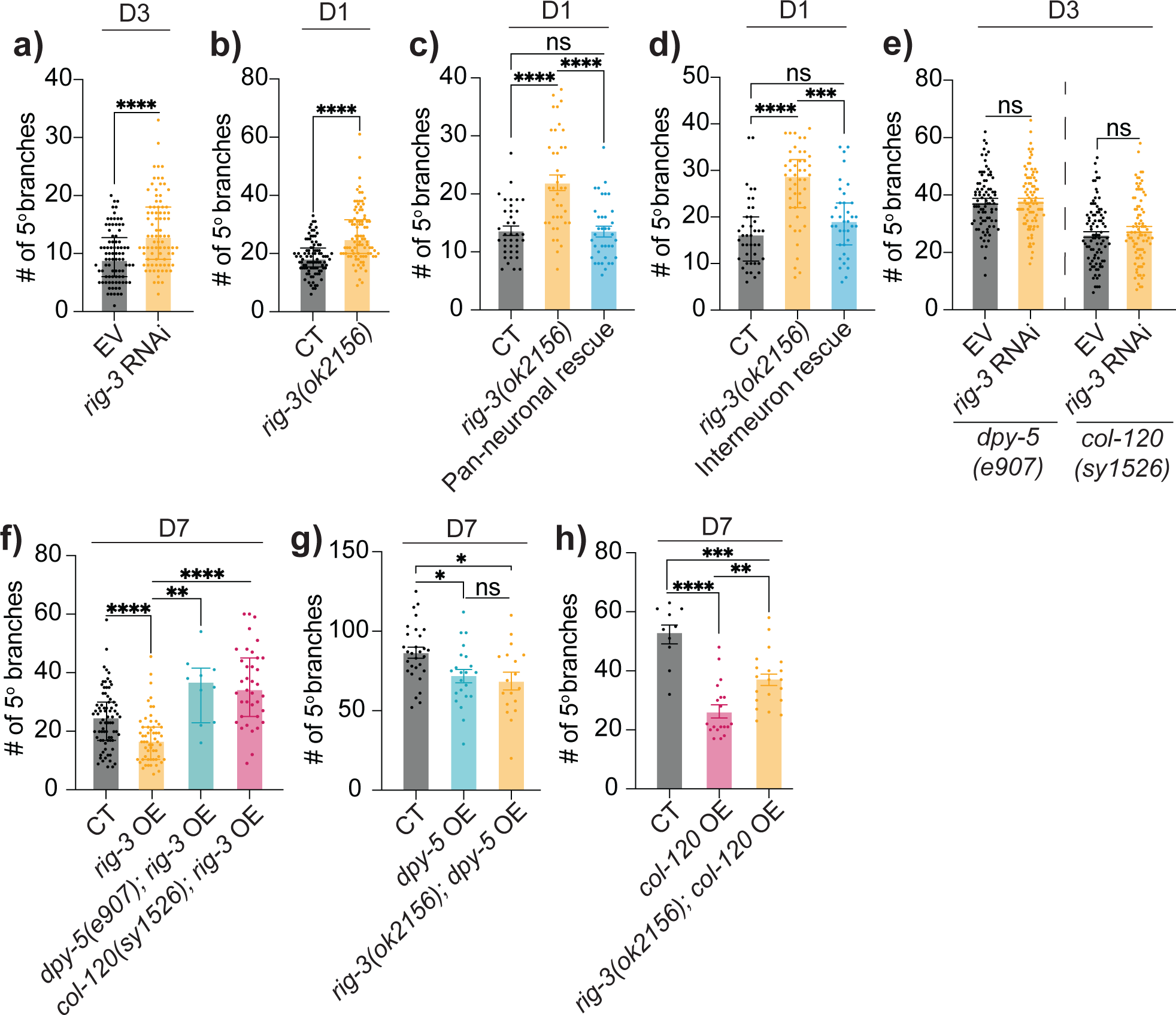
RIG-3/IgSF acts as upstream of epidermal collagens to regulate PVD dendritic integrity in aging. **(a)** Quantification of 5° dendrites in empty vector (EV) and *rig-3* RNAi treated CT animals at D3. (n = 84). **(b)** 5° dendrites in CT (n = 91) (*ser-2(3)p::GFP(lxyEx77)*, see Methods) and *rig-3(ok2156)* mutant (n = 90). **(c)** 5° dendrites in CT (n = 37), *rig-3(ok2156)* (n = 40), and pan-neuronal rescue (driven by *unc-33p*) in *rig-3(ok2156)* (n = 37). **(d)** 5° dendrites in CT (n = 41), *rig-3(ok2156)* (n = 46), and interneuron rescue (driven by *nmr-1p*) in *rig-3(ok2156)* (n = 37). **(e)** 5° dendrites in *dpy-5(e907)* mutant animals treated with EV (n = 84) or *rig-3* RNAi (n = 85), as well as in *col-120(sy1526)* mutant animals treated with EV (n = 88) or *rig-3* RNAi (n = 86). **(f)** 5° dendrites in CT (n = 76), *rig-3* pan-neuronal overexpression driven by *unc-33p* (n = 57), *rig-3* overexpression in *dpy-5(e907)* mutant background (n = 10), and *rig-3* overexpression in *col-120(sy1526)* mutant background (n = 37). **(g)** 5° dendrites in CT (*ser-2(3)p::GFP(lxyEx77)*) (n = 30), *dpy-5* epidermis-specific overexpression driven by *col-19p* (n = 22), and *dpy-5* overexpression in *rig-3(ok2156)* mutant background (n = 18). **(h)** 5° dendrites in CT (n = 10), *col-120* epidermis-specific overexpression driven by *col-19p* (n = 19; line 1), and *col-120* overexpression (line 1) in *rig-3(ok2156)* mutant background (n = 20). ns-not significant, * *p*<0.05, ** *p*<0.005, *** *p*<0.001, **** *p*<0.0001.

We then conducted a rescue experiment using a pan-neuronal promoter to drive wild-type *rig-3* expression in *rig-3(ok2156)*, fully restoring normal PVD dendritic branching (Figure 6c, Figure S7g). Pan-neuronal overexpression of *rig-3* also significantly suppressed excessive branching in aged animals (Figure 6f), similar to the effects of *col-120* or *dpy-5* overexpression (Figure 4). Since *rig-3* is not detected in PVD, we sought to identify the specific neuron(s) through which it acts. *rig-3* is highly expressed in interneurons, particularly in AVA, where it regulates locomotion by modulating neuropeptide release from AVA, which in turn influences sensory neurons connected to AVA (Bhardwaj et al., 2020). Notably, PVD axon also synapses with AVA (White et al., 1986), leading us to hypothesize that RIG-3 may regulate PVD dendritic integrity from AVA. Using the *nmr-1* promoter, which is active in fewer than 10 neuron subsets— primarily interneurons including AVA, AVD, AVE and PVC, all of which also endogenously express *rig-3* (Figure. S7i) (Brockie et al., 2001; Taylor et al., 2021), we conducted a rescue experiment in *rig-3(ok2156)*. This fully rescued the excessive branching phenotype, suggesting *rig-3* non-cell-autonomously regulates PVD dendritic branching from within this small set of interneurons.

To determine if *rig-3* mediates the effects of skin collagens on PVD branching, we performed adulthood RNAi of *rig-3* in *col-120* or *dpy-5* mutants and found that it did not further enhance excessive branching in collagen mutants, suggesting *rig-3* functions in the same pathway as *dpy-5* and *col-120* in regulating PVD dendritic integrity during aging (Figure 6e, Figure S7j). Further epistasis analysis surprisingly revealed that *rig-3* overexpression’s beneficial effect on branching in aged animals was completely blocked by *col-120* or *dpy-5* mutations (Figure 6f, Figure S7k). Conversely, *col-120* or *dpy-5* overexpression’s effects remained even with *rig-3* mutation (Figure 6g-h, Figure S7l-m). Contrary to our expectations, these data suggest that neuronal RIG-3 likely functions upstream of collagens in maintaining PVD dendritic health over the course of aging.

### 2.9 Beneficial effect of COL-120 on PVD dendritic integrity requires DAF-16/FOXO

To further explore downstream pathways of skin collagen genes, we analyzed transcriptomic data (GSE19310) comparing wild-type animals and *dpy-10(e128)* mutants, another cuticular collagen mutation associated with early-onset excessive PVD branching (Figure S4). KEGG pathway analysis of differentially expressed genes in *dpy-10(e128)* revealed significant downregulation in several signaling pathways, including FOXO signaling (Figure S7n). We sought to examine potential interactions between collagen genes and *daf-16/FOXO*, focusing on *col-120*, given its stronger effect in suppressing excessive branching when overexpressed (Figure 4). We found that *daf-16(mu86)* mutation completely suppressed the beneficial effect of epidermis-specific *col-120* overexpression (Figure S7o-p). Since *col-120* overexpression was driven by the *col-19* promoter specifically in adulthood, rather than by its endogenous promoter, it is unlikely that the observed effect involves *daf-16* regulation of *col-120* expression. Additionally, previous studies show that *col-19* transcription is unaffected by *daf-2* or *daf-16* (Ewald et al., 2015; Uno et al., 2013), indicating that the effect observed with *daf-16* mutation is unlikely due to reduced expression of *col-19p::col-120* either. Notably, the suppression effect caused by *daf-16* mutation in *col-120* overexpression animals does not appear to result from the reduced lifespan caused by *daf-16* mutation. This interpretation is supported by the observation that D7 animals with *daf-16(mu86)* and *col-120* overexpression displayed branching levels more comparable to age-matched CTs, unlike the significant difference observed between D7 CT and D7 *daf-16(mu86)* single mutants (Figure 1h, Figure S1g). Together, these results suggest that *daf-16*/FOXO may act downstream of *col-120* in regulating PVD dendritic branching during aging.

## 3. DISCUSSION

Our study demonstrates that age-related reduction in epidermal collagen drives both structural and functional changes of sensory neuron aging process. Interestingly, the impacts exhibit selectivity, manifested at two levels: firstly, collagen reduction does not contribute to neuritic beading in aging neurons, pointing to a phenotype-specific effect; secondly, their role is confined to regulating branching integrity in specific sub-types of cutaneous sensory neurons, highlighting a cell-type-targeted manner in neuron-epidermal interactions. Combined with our previous findings (E et al., 2018), these results suggest that various molecular cues from non-neural tissues during aging may trigger distinct neuronal aging mechanisms. Given that the investigated collagens are secreted to the apical surface of the epidermis, their influence on sensory neuron aging likely occurs via indirect mechanisms, rather than direct structural support for neuronal structures. Unexpectedly, we identify RIG-3/IgSF in interneurons as an upstream partner of skin collagens, coordinating neuronal homeostasis during aging. Notably, the protective effect of collagen overexpression requires DAF-16/FOXO, but independent of its role in lifespan regulation. Taken together, our findings suggest that the regulatory role of skin collagens is mediated through an intricate, bi-directional communication processes between neurons and epidermal cells, emphasizing the need for a paradigm shift towards exploring multi-tissue strategies to comprehensively address the complexities of neuronal aging.

Although age-related declines in collagen production in various tissues have been well-documented, its impact on the nervous system during aging remains underexplored. One study reported that aged mice lacking COL6A1, a ubiquitously expressed collagen VI gene, exhibited more severe deficits in motor coordination and spatial memory compared to age-matched WT animals (Cescon et al., 2016). Our current study extends this area by demonstrating that non-basement membrane collagens, particularly those from non-neural tissues, play a distinctive regulatory role in maintaining neuronal structure and function during aging. While our data suggest that DAF-16/FOXO mediates COL-120’s effects on PVD dendritic health, questions remain about whether DAF-16 acts within epiderma cells, whether COL-120 engages additional signaling pathways that interact with DAF-16 elsewhere, and which specific DAF-16 target genes are involved. Importantly, changes in key collagen genes like *col-120* could broadly alter extracellular matrix (ECM) composition (Teuscher et al., 2024) or influence signaling pathways, such as Wnt and TGF-β, as seen with *dpy-10* (Figure S7). In fact, such extensive downstream effects may complicate the rescue of PVD excessive branching phenotype in *col-120* or *dpy-5* mutants, as observed in this study, even with endogenous promoters, as compensatory responses induced by collagen mutations could interfere with attempts to fully restore normal function.

An unexpected finding from our study is the role of RIG-3/IgSF in interneurons as an upstream partner of skin collagens in modulating PVD dendritic branching. Our data suggest that RIG-3 acts from interneurons, likely adjacent to PVD, yet the beneficial effects of *rig-3* overexpression are blocked by the loss of skin collagens. This observation implies a signaling loop between neurons and epidermal cells, rather than a simple neuron-neuron interaction, in the regulation of neuronal aging. RIG-3 shares homology with mammalian neural cell adhesion molecule 1 (NCAM1), also an IgSF protein (Babu et al., 2011). Studies in a rat sciatic nerve injury model have shown that NCAM1 on axonal membrane is essential for axonal fasciculation during nerve regeneration, requiring interaction between extracellular collagen VI and the fibronectin type III (FNIII) domain of NCAM1 (Sun et al., 2022). While RIG-3 also contains a FNIII domain (Babu et al., 2011), the segregation of cuticular collagens to the epidermal apical surface makes direct interaction with COL-120 or DPY-5 unlikely. Notably, RIG-3 can function as either a cell surface or secreted protein (Babu et al., 2011); future studies may clarify whether its secretion is required for its role in regulating PVD branching and assess its potential to act over long distances.

Excessive branching in aging peripheral neurons is a conserved phenomenon. For example, elderly humans with sensory neuropathy exhibit increased branching of epidermal nerve fibers in the proximal thigh compared to healthy controls (Lauria et al., 1999). Our time-lapse imaging revealed that excessive high-order dendritic branches in PVD neurons are enriched with F-actin but lack microtubules, indicating high structural plasticity, much like dendritic spines, known to affect neuronal signaling and synaptic plasticity. Therefore, it is tempting to speculate that increased branching observed in aging reflects a compensatory plasticity for future neurodegeneration at more advanced ages. Mechanotransduction may play a role in this phenomenon. As *C. elegans* age, their cuticle becomes stiffer, while the ECM weakens, reducing resistance to mechanical stress. Studies have shown that *col-120* expression can be upregulated in response to mild mechanical compression, and loss of *col-120* decreases cuticle mechanical resistance, particularly in long-lived *daf-2* mutants, suggesting a feedback loop between ECM homeostasis and mechanotransduction (Rahimi et al., 2022; Teuscher et al., 2024). Mechanosensitive sodium channels, such as MEC-10, UNC-8, DEL-1, and DEGT-1, are localized to PVD dendritic branches and are essential for proprioceptive and mechanonociceptive signaling (Tao et al., 2019). Age-related collagen loss could compromise ECM stability, allowing cumulative mechanical stress to alter channel sensitivity over time and, in turn, influence dendritic branching through modified sensory feedback to mechanical cues. This effect may ultimately contribute to compensatory branching, inadvertently engaging sensorimotor networks beyond their usual functional scope.

During nervous system development, transmembrane and extracellular proteins are crucial for shaping neuronal structure and guiding migration. Proteins such as SAX-7/L1CAM, MNR-1, and DMA-1 (transmembrane LRR protein) have been extensively studied for their roles in PVD dendritic morphogenesis; however, deficiencies in these proteins primarily affect 1°-4° dendritic branches and do not contribute to excessive higher-order branching (Dong et al., 2013; Salzberg et al., 2013). Additionally, we did not observe a clear link between abnormal dendrite overlapping, often due to self-avoidance defects (Kravtsov et al., 2017; Oren-Suissa et al., 2010), and excessive branching either during aging or in collagen mutants. These observations suggest that the molecular mechanisms responsible for neuron morphogenesis in development may be distinct from those driving neuron structural deterioration in aging.

Our findings indicate that polymodal sensory neurons can undergo multiple, distinct morphological changes during aging, each corresponding to specific functional declines. In PVD neurons, simultaneous excessive dendritic branching and dendritic beading are rare, suggesting that animals may avoid experiencing major declines in both harsh touch function and proprioception concurrently. The current study combined with our previous research (E et al., 2018) indicate that within multifunctional neurons, certain structures and functions may be selectively vulnerable to aging, likely driven by distinct non-neural signals. The question of whether polymodal neurons prioritize one function over another remains unanswered.

On a broader scale, neurons across different regions exhibit varying susceptibility or resistance to aging, leading to differences in the onset timing and progression of neurodegenerative processes. This selective vulnerability is commonly observed in both normal aging and age-related neurodegenerative diseases, and evident across species, including *C. elegans*. For instance, ALM, PLM, and PVD neurons age at different rates (E et al., 2018) and respond differently to the same skin-derived aging cues. Given that each neuron class/sub-type has a unique gene expression profile, future research should investigate whether these molecular signatures dictate responsiveness to non-neural or extracellular aging cues, contributing to selective vulnerability.

## 4. MATERIALS AND METHODS

### 4.1 *C. elegans* Strains and Maintenance

*C. elegans* strains were maintained at 20°C, or as otherwise indicated, on Nematode Growth Medium (NGM) plates seeded with *E. coli* OP50. Only hermaphrodites were used for data generating experiments. For age-synchronized progeny, plates with gravid hermaphrodites were bleached to eliminate microbes, and animals were transferred to NGM plates containing 100 μM 5’ fluorodeoxyuridine (FUDR) (VWR, 76345-984) at the mid-or late-L4 stage to prevent egg hatching. Day 1 of adulthood (D1) was defined as 24 hours post-L4. Given the potential confounding effects of FUDR on aging (Feldman et al., 2014; Van Raamsdonk & Hekimi, 2011), key experiments were repeated without FUDR, moving animals onto fresh plates daily until D5, every two days thereafter. FUDR-free experiments are noted in figure legends. See Table S3 for strain information.

### 4.2 DNA Constructs and Generation of Transgenes

DNA expression constructs were generated using Gateway cloning technology (ThermoFisher), with details in Table S4. Primers were acquired from Millipore and ThermoFisher (Table S5). Transgenic animals were generated by microinjection, with plasmid DNAs used at 10-100 ng/μL, co-injection marker *ttx-3p::RFP* or *ttx-3p::GFP* at 50 ng/μL (Tables S3, S4). At least two independent transgenic lines were analyzed per construct. Promoters used include—*dpy-5p* (-854 bp from start codon), *col-120p* (-348 bp), *nmr-1p* (-1104 bp), *dpy-7p* (-304 bp), *col-19p* (-649 bp), and *unc-33p* (-1974 bp).

### 4.3 Fluorescent Microscopy

Neurons were visualized using fluorescent reporters and phenotyped on a Zeiss Axio Imager M2 using 0.5-1% M9 dilution of 1-phenoxy-2-propanol (Fisher) for immobilization. Confocal images were taken on a Leica SP8 with Z-stack (0.5-1µm/slice) at 63X magnification, using polystyrene beads (Polysciences, 00876-15) for immobilization.

### 4.4 Quantification of Neuron Morphological Defects

*wdIs51(F49H12.4::GFP)* (Smith et al., 2010) was used to visualize PVD neurons. Excessive dendritic branching was quantified by counting 5° and 6° branches, identified within ‘menorah’ structures (Figure 1b). 5° and 6° branches for each animal were counted for either PVDL or PVDR for either the dorsal or ventral side, depending on which was clearly visible during the imaging session. PVD dendritic beading phenotype was assessed by counting bead- or bubble-like structures along the dendrites of either PVDL or PVDR across the entire dendritic tree. Animals with ≥ 10 beads across ≥ 5 menorahs were considered positive for degeneration (E et al., 2018). Severity scoring: in Figure 5a, animals with 0-10 beading structures was scored as normal, 10-20 as mild, 20-50 as moderate, and >50 as severe; in Figure S1b, a finer scale was used: score 1 for 0-10 structures, 2 for 10-20, 3 for 20-30, 4 for 30-50, 5 for 50-100, 6 for 100-200, and 7 for over 200. *ser-2(3)p::mCherry-PH* was used to visualize PVD dendritic membrane structures with respect to F-actin dynamics. For *rig-3* mutants, *ser-2(3)p::GFP (lxyEx77)* was constructed to visualize PVD neurons due to challenges in crossing *rig-3(ok2156)X* with *wdIs51X*. *ser-2(3)p::GFP(lxyEx77)* animals had larger body size than *wdIs51*, likely due to genetic background differences, which may account for the branching number variation at D1 between the two reporter strains.

*zdIs5(mec-4p::GFP)* was used to visualize ALM and PLM neurons, and the number of ectopic branches sprouting from their main neurites were counted.

For each experimental group, a minimum of three replicates of 30 animals, or indicated otherwise, were quantified alongside a control group, with experimenters blinded to sample identity to minimize bias.

### 4.5 PVD Function Analysis: Harsh Touch Response

The measurement of the harsh touch response was conducted using a *mec-4(u253)* mutant background with non-functional touch receptor neurons but intact PVD neuron functionality, as previously described (E et al., 2018), with modifications to the scoring system. A score of 1 was given if the animal moved over one body length, 0.5 if it moved between one-third and one body length, and 0 if it moved less than one-third or not at all. Each animal underwent 10 trials with 10-15 minute intervals. Cumulative scores ranged from 0 (severely impaired response) to 10 (optimal response).

### 4.6 PVD Function Analysis: Proprioception

Proprioceptive sensing specific to PVD neurons was assessed using a locomotion assay as described previously (Tao et al., 2019; Tavernarakis et al., 1997) with some modifications. On the day of assay, animals were transferred to fresh NGM plates seeded with OP50, one animal per plate, and allowed to roam freely for 2-3 hours at 20°C, until sufficient tracks were made in the bacterial lawn. The plate was then imaged using a digital camera attached to a Leica S9i stereomicroscope at 1X magnification.

Three independent replicates of 10 animals each were assayed for each group. A total of 100 randomly selected wavelengths (distance between two adjacent peaks) and amplitudes (distance between peak and wavelength, perpendicular to wavelength) were measured using Fiji. All measurements were used in cases where animals had fewer than 100 measurements, with a minimum of 40 measurements per animal. All measurements were normalized to the body lengths of individual animals and units were converted to millimeters utilizing a reference slide. All animals subjected to proprioception assay were also examined under the Zeiss to quantify PVD dendritic branching.

### 4.7 RT-qPCR

To quantify the change in expression level of collagens in aging, N2 WT animals at D1 and D7 were collected, washed with M9 solution for three 30-minute washes, and total RNA was isolated with TRI reagent (Millipore Sigma).

To test RNAi efficacy, 10-worm lysis method (Ly et al., 2015) was used to prepare lysates. Briefly, 10 worms cleaned in H_2_O and placed in 1 µL of lysis buffer (0.25 mM EDTA, 5mM Tris pH 8., 0.5% Triton X-100, 0.5% Tween-20, 1mg/mL proteinase K). Samples were lysed in a thermal cycler at 65°C for 15 minutes, 85°C for 1 minute, and stored at -80°C for at least 12 hours. An additional 1.5 µL of lysis buffer was then added, and the cycle was repeated. Samples were treated with dsDNAse (ThermoFisher; 0.5 µL each of 10X buffer and dsDNAse, 1.5 µL ultrapure H2O) and incubated at 37°C for 15 minutes, 55°C for 5 minutes, and 65°C for 1 minute, followed by reverse transcription.

Reverse transcription was performed with iScript Reverse Transcription Supermix for RT-qPCR (BioRad). RT-qPCR was performed using either TaqMan Gene Expression Assays (ThermoFisher, Ce02462726_m1, Ce02417736_s1, Ce02457574_g1) and iTaq Universal Probes Supermix (Bio-Rad) or using designed primers (Table S5) and iTaq Universal SYBR Green (Bio-Rad). *ama-1* and *cdc-42* were used as housekeeping controls. Primer efficiency was verified utilizing standard curve method. Bio Rad CFX96 Touch Deep Well Real-time PCR Detection System was used to perform the qPCR and relative mRNA levels were calculated using comparative ΔΔCT method.

### 4.8 Lifespan Assay

L4 animals were transferred to assay plates and scored daily. An animal was considered dead when it no longer responded to gentle prodding with a platinum wire and displayed no pharyngeal pumping. Lifespan was calculated from the time animals were first placed on the assay plates until they were scored as dead. Animals that died due to protruding/bursting vulva, internal hatching, as well as animals that crawled off the agar were censored. For lifespan assays examining collagen overexpression strains, the PVD::GFP background strain (*wdIs51*) was used as the control.

### 4.9 Time-lapse Imaging

Imaging of PVD branches was done on the Leica SP8 using 2% agarose pads with polystyrene beads for immobilization. Control animals *[ser2(3)p::mCherry-PH; ser2(3)p::GFP::UtrCH(lxyEx51)]* were imaged at D1 and D9. Each session lasted 8 minutes with 2-minute intervals, and images were Z-stack projections captured with a 63X objective in both red and green channels. Branch dynamics were quantified in Fiji by measuring the length of individual higher-order branches at each time-point, and actin dynamics were assessed by observing GFP-labeled actin movement.

### 4.10 Adulthood-specific RNAi

Feeding method was used to perform RNAi experiments (Conte et al., 2015). FUDR at a concentration of 100 µM was externally added to RNAi plates following growth of the HT115 bacterial lawn. The RNAi clones were obtained from the Vidal RNAi library (Horizon Discovery, Table S4) or amplified from cDNA using PCR. The section of cDNA included for each RNAi clone are *dpy-5::L4440 (1)* (PELZ88: 210 bp section downstream of *dpy-5* start codon) and *col-120::L4440 (1)* (PELZ45: 256 bp section downstream of *col-120* start codon). These regions were selected to ensure RNAi specificity, confirmed by BLAST search to ensure no homology with other collagen genes. Additional RNAi clones were also included: *dpy-5::L4440 (2)* (PELZ48: 554 bp section 301 bp downstream of *dpy-5* start codon) and *col-120::L4440 (2)* (PELZ46: 605 bp section 337 bp downstream of *col-120* start codon). All RNAi clones were sequenced to verify their identities. Empty L4440 plasmid was used as a negative control. TJ356 *[zIs356 (daf-16p::daf-16a/b::GFP)]* animals with empty-vector or *daf-2* RNAi treatment were used as a positive control: Successful RNAi condition was indicated by nuclear translocation of DAF-16::GFP in *daf-2* RNAi-treated animals, while cytoplasmic localization indicated unsuccessful treatment.

### 4.11 Gene expression datasets and pathway analysis

Gene expression datasets (GSE176088, GSE19310) were obtained from the NCBI Gene Expression Omnibus. Data were analyzed using GEO2R to identify differentially expressed genes (DEGs), with criteria of log2-fold change ≥ 1.5 and a *p*-value < 0.05. KEGG enrichment analysis was performed using the Database for Annotation, Visualization and Integrated Discovery (Sherman et al., 2022), applying an FDR < 0.01 for statistical significance. The top 10 significant KEGG pathways were visualized in a bubble graph, with the rich factor representing the proportion of DEGs within each pathway term.

### 4.12 Statistical Analysis

Data processing, statistical analyses, and graph plotting were performed with GraphPad Prism software. Normality was assessed using the D’Agostino & Pearson test. For comparisons between experimental and control groups, unpaired t-test for normally distributed data, and the Mann-Whitney test for non-normally distributed data were used. When comparing three or more groups, either one-way ANOVA or the Kruskal-Wallis test, followed by post-hoc analysis. Pearson’s correlation test was applied for correlation analyses. Log-Rank test was used to analyze lifespan data. Bar graphs display the mean ± SEM for normally distributed data, and the median with interquartile range for non-normally distributed data.

## ACKNOWLEDGMENTS

We thank Bradey A. R. Stuart for assistance in molecular cloning and construction of transgenic strains. Some strains were provided by the CGC, which is funded by NIH Office of Research Infrastructure Programs (P40 OD010440).

## CONFLICT OF INTEREST

The authors declare that the research was conducted in the absence of any commercial or financial relationships that could be construed as a potential conflict of interest.

## FUNDING SUPPORT

This work was supported by the Imagine More Award and Research Affairs Committee at the Medical College of Wisconsin, as well as Advancing a Healthier Wisconsin (AHW) Endowment project 5520482 titled Developing Innovative Translational Research Programs in Clinically Relevant Neurological Disorders.

## AUTHOR CONTRIBUTIONS

L.E. conceived the project; L.E. and M.K. designed the experiments; M.K., L.E., S.G.W., A.L.F., and E.M. performed the experiments and collected the data; M.K., S.G.W. and L.E. analyzed and interpreted the data; M.K. and L.E. drafted the manuscript; M.K., S.G.W., A.L.F., E.M. and L.E. edited and proofread the manuscript.

## DATA AVAILABILITY

The data that support the findings of this study are available from the corresponding author upon reasonable request.

## Supplemental Materials

### SUPPLEMENTAL FIGURE LEGENDS

**Figure S1:**
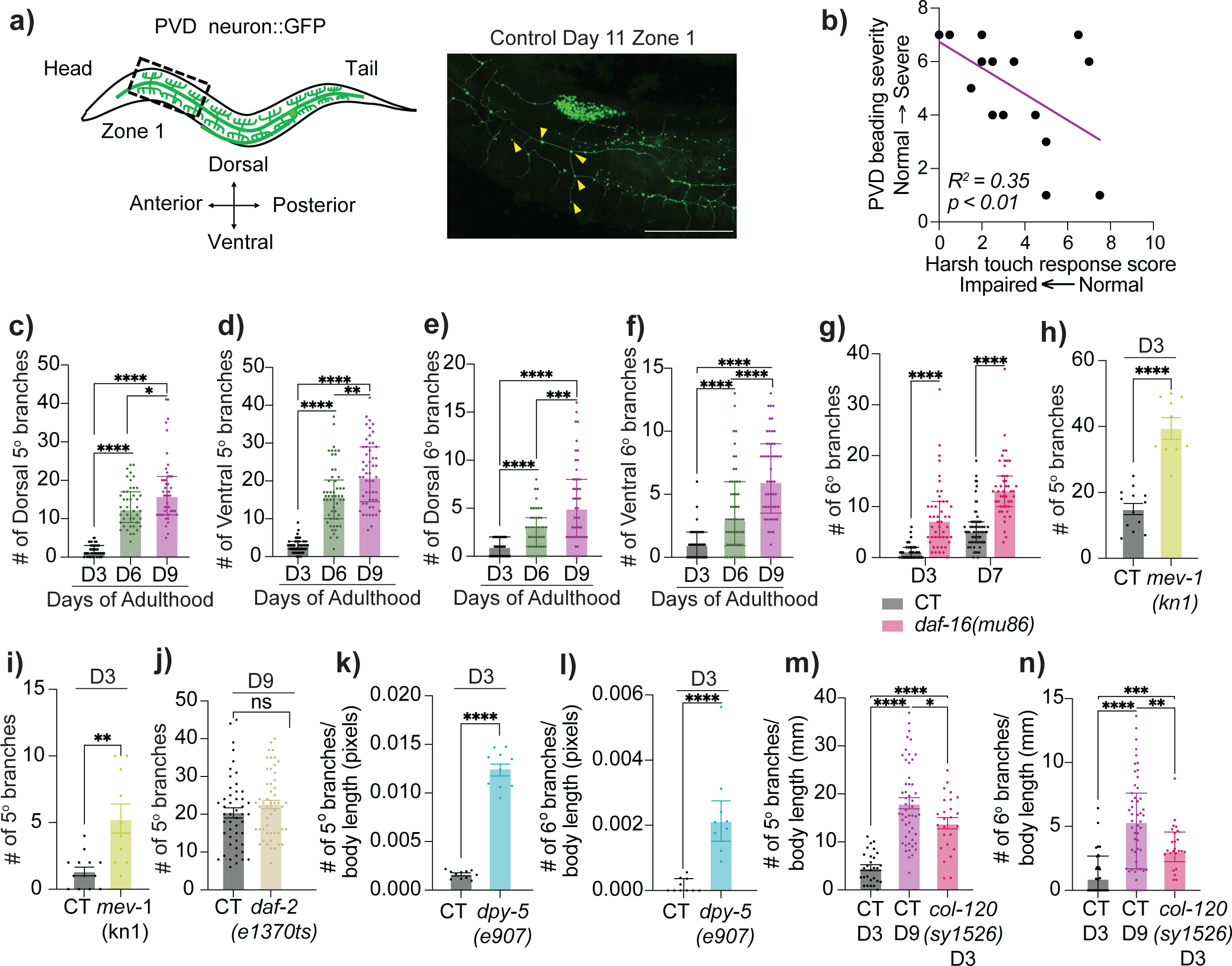
Characterization of the aging PVD neuron. **(a)** Schematic diagram of the PVD neuron in control (CT) animals. The black box indicates the zone presented in the representative image of CT animals at Day 11 (D11) of adulthood. Yellow arrowheads indicate neuritic beads. Scale bar = 50 μM. **(b)** Correlation between PVD beading severity and harsh touch response in CT [*mec-4(u253)*; *F49H12.4::GFP(wdIs51)*] animals at D7. See Methods for scoring criteria. *n* = 18. **(c-f)** Number of dorsal *(c)* and ventral *(d)* 5° dendrites, and dorsal *(e)* and ventral *(f)* 6° dendrites in CT [*F49H12.4::GFP(wdIs51)*] animals at D3 (Dorsal: n = 50, Ventral: n = 55), D6 (Dorsal: n = 44, Ventral: n = 54), and D9 (Dorsal: n = 47, Ventral: n = 53). **(g)** 6° dendrites in CT and *daf-16(mu86)* mutant animals at D3 and D7 (CT D3: n = 51, CT D7: n = 70, *daf-16* mutant D3: n = 47, *daf-16* mutant D7: n = 46). Experiments performed without FUDR. **(h-i)** 5° *(h)* and 6° *(i)* dendrites in CT (n = 13) and *mev-1(kn1)* (n = 10). Experiments performed without FUDR. **(j)** 5° dendrites in CT (n = 49) and *daf-2(e1370)* (n = 51). **(k-l)** 5° *(k)* and 6° *(l)* dendrites normalized to body length in CT (n = 11) and *dpy-5(e907)* (n = 10). **(m-n)** 5° *(m)* and 6° *(n)* dendrites normalized to body length in CT (D3: n = 28, D9: n = 51) and *col-120(sy1526)* (n = 25). ns-not significant, * *p*<0.05, ** *p*<0.005, *** *p*<0.001, **** *p*<0.0001.

**Figure S2:**
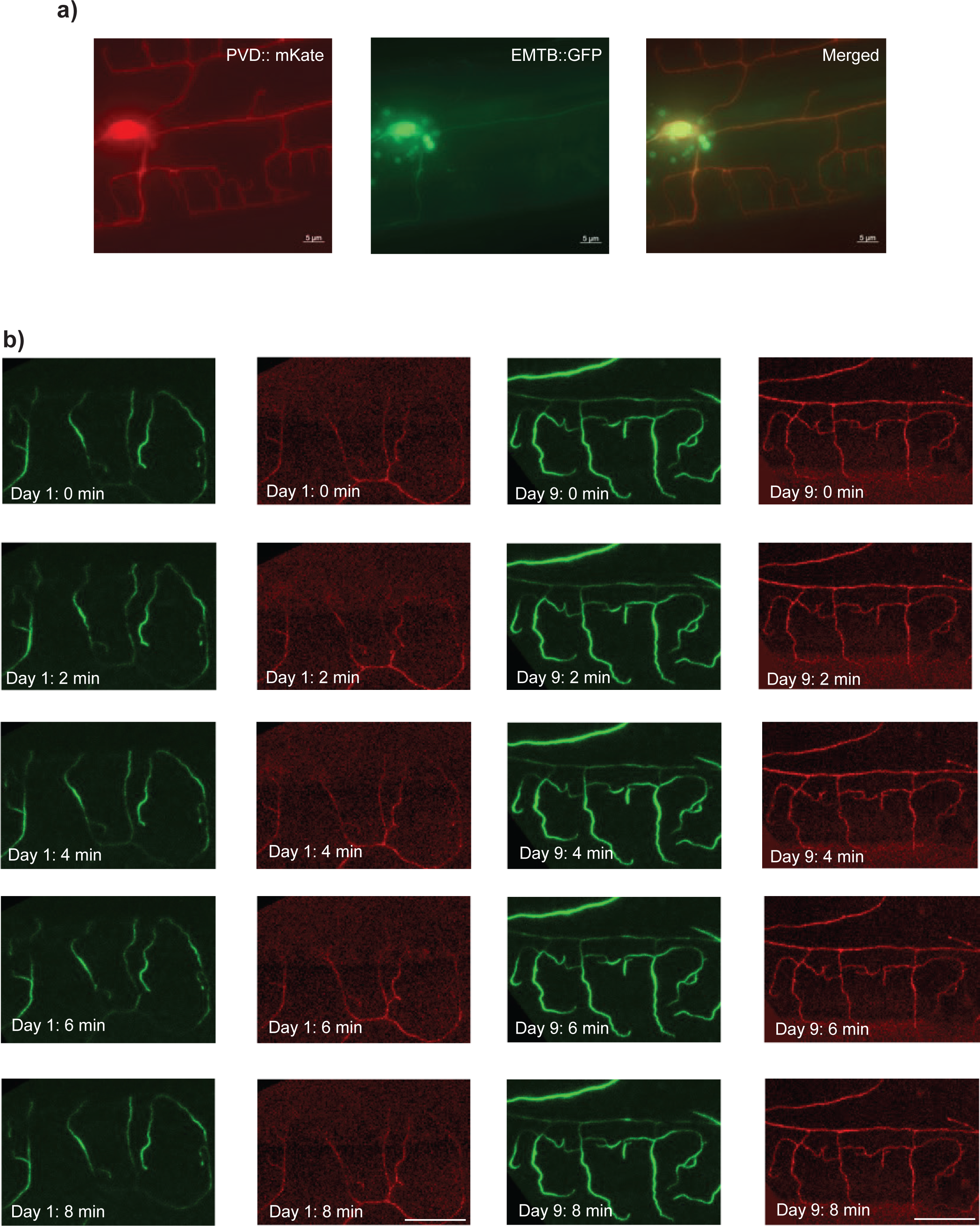
Cytoskeletal composition in PVD dendrites. **(a)** Split channel and merged images of microtubule localization in the PVD neuron at D7 in control (CT) animals [*ser-2(3)p::EMTB::GFP + ser-2(3)p::mCherry-PH(lxyEx48)*]. Red channel displays the PVD neuron, while the green channel displays the microtubules in the PVD. **(b)** Split channel representative time-lapse images of actin dynamics in higher order dendrites during an 8-minute session at 2-minute intervals at D1 and D9 of adulthood in CT animals [*ser-2(3)p::GFP::UtrCH + ser-2(3)p::mCherry-PH(lxyEx51)*]. Red channel displays the PVD neuron, while the green channel displays the actin in the PVD.

**Figure S3:**
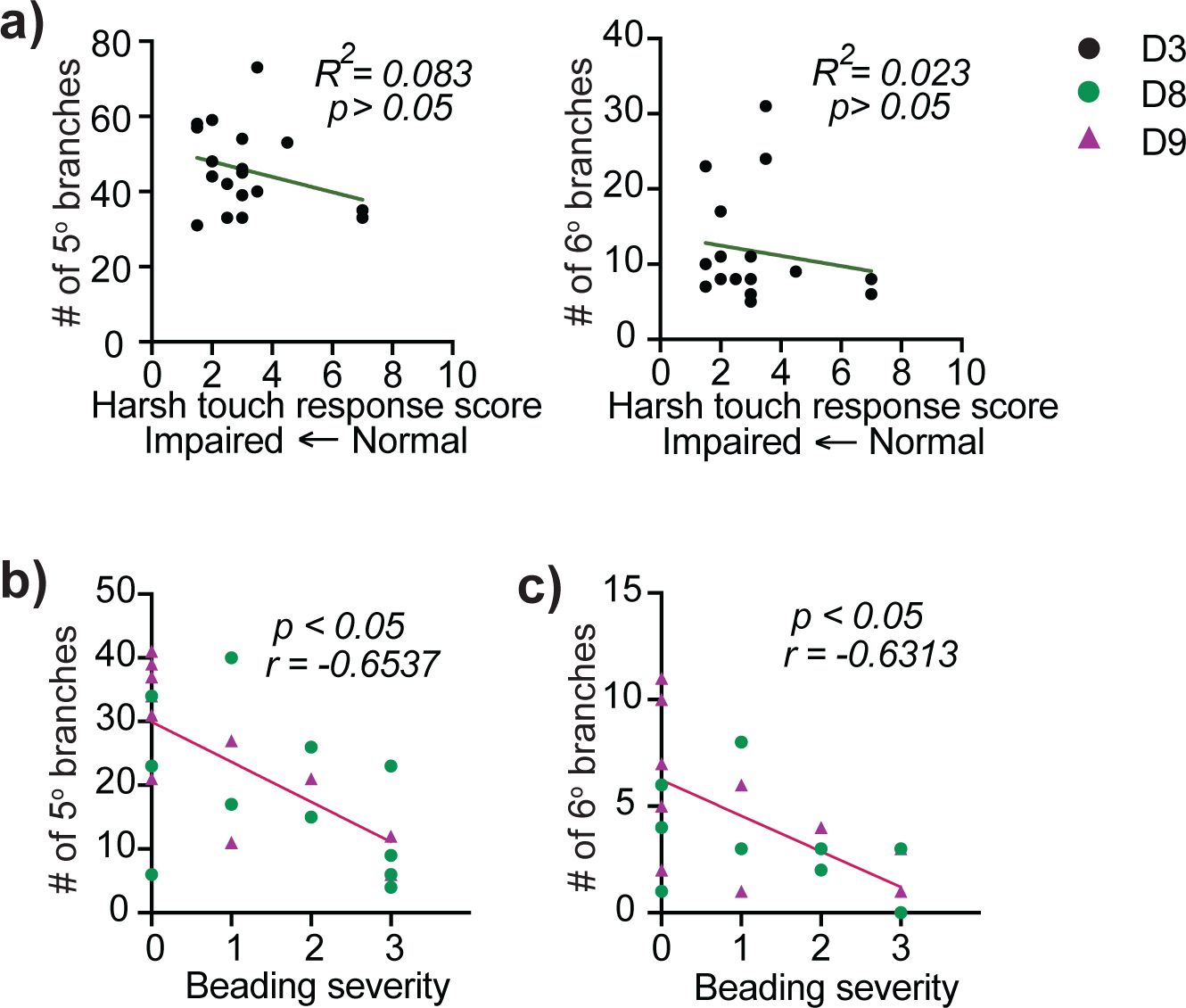
Aging-associated dendritic branching correlated with proprioceptive deficits and neuritic beading. **(a)** Correlation between number of higher-order dendrites (5° and 6°) and harsh touch response in *mec-4(u253)* mutant animals at D7. Harsh touch response was scored based on the distance moved with a cumulative score ranging from 0 to 10, indicating impaired to normal response respectively. n = 18. **(b-c)** Correlation between higher-order branching and the extent of neuritic beading in CT animals at D8 (n = 11) and D9 (n = 11). Higher-order dendrites are 5° dendrites *(b)* and 6° dendrites *(c)*. Beading severity is measured as 0-normal, 1-mild, 2-moderate, and 3-severe.

**Figure S4:**
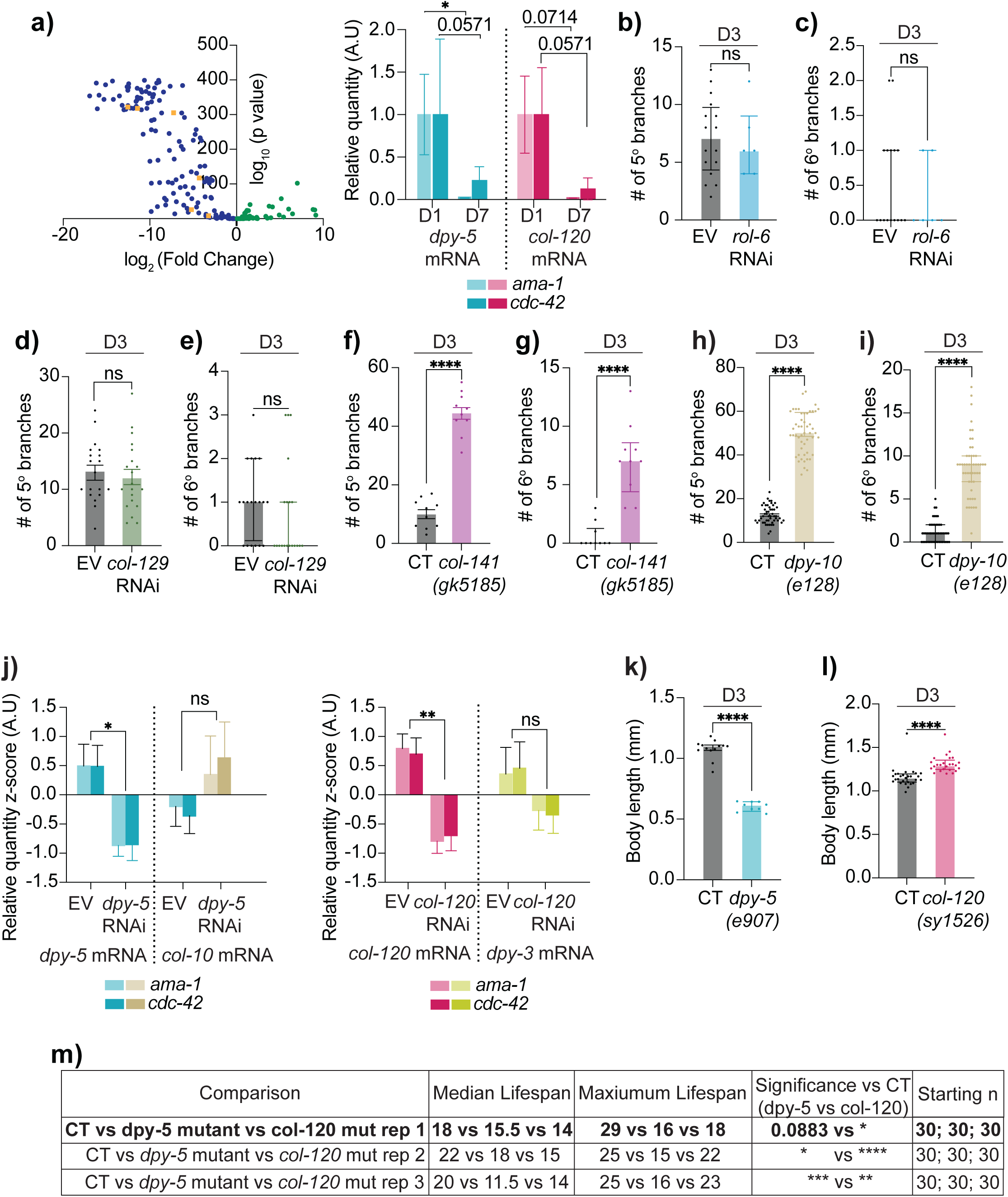
Characterization of cuticular collagens in aging. **(a)** (Left) Volcano plot depicting changes in expression of skin collagens in D10 animals relative to L4 animals from the GSE176088 dataset. *col-120, col-129, col-141, dpy-5, dpy-10, and rol-6* are shown in orange. (Right) qPCR verification of *dpy-5* and *col-120* mRNA expression levels in CT animals at D1 and D7, represented as relative quantity, with *ama-1* or *cdc-42* as housekeeping genes. **(b-c)** Quantification of 5° *(b)* and 6° *(c)* dendrites in empty vector (EV) (n = 16) and *rol-6* RNAi (n = 6). **(d-e)** 5° *(d)* and 6° *(e)* dendrites in EV (n = 19) and *col-129* RNAi (n = 20). **(f-g)** 5° *(f)* and 6° *(g)* dendrites in CT and *col-141(gk5185)* mutant (n = 10). **(h-i)** 5° *(h)* and 6° *(i)* dendrites in CT and *dpy-10(e128)* mutant (*n* = 50). **(j)** mRNA levels of *dpy-5* and *col-10* (predicted paralog of *dpy-5*) after *dpy-5* RNAi clone 1 treatment *(left),* and mRNA levels of *col-120* and *dpy-3* (predicted paralog of *col-120*) after *col-120* RNAi clone 1 treatment *(right)*, relative to empty vector (EV), with *ama-1* or *cdc-42* as housekeeping genes. **(k)** Comparison of body lengths between control (CT, n = 13) and *dpy-5(e907)* (n = 9). **(l)** Comparison of body lengths between control (CT, n = 28) and *col-120(sy1526)* (n = 25). **(m)** Summary data of lifespan analysis of CT, *dpy-5* mutant and *col-120* mutant animals. ns- not significant, * *p*<0.05, ** *p*<0.005, *** *p*<0.001, **** *p*<0.0001.

**Figure S5:**
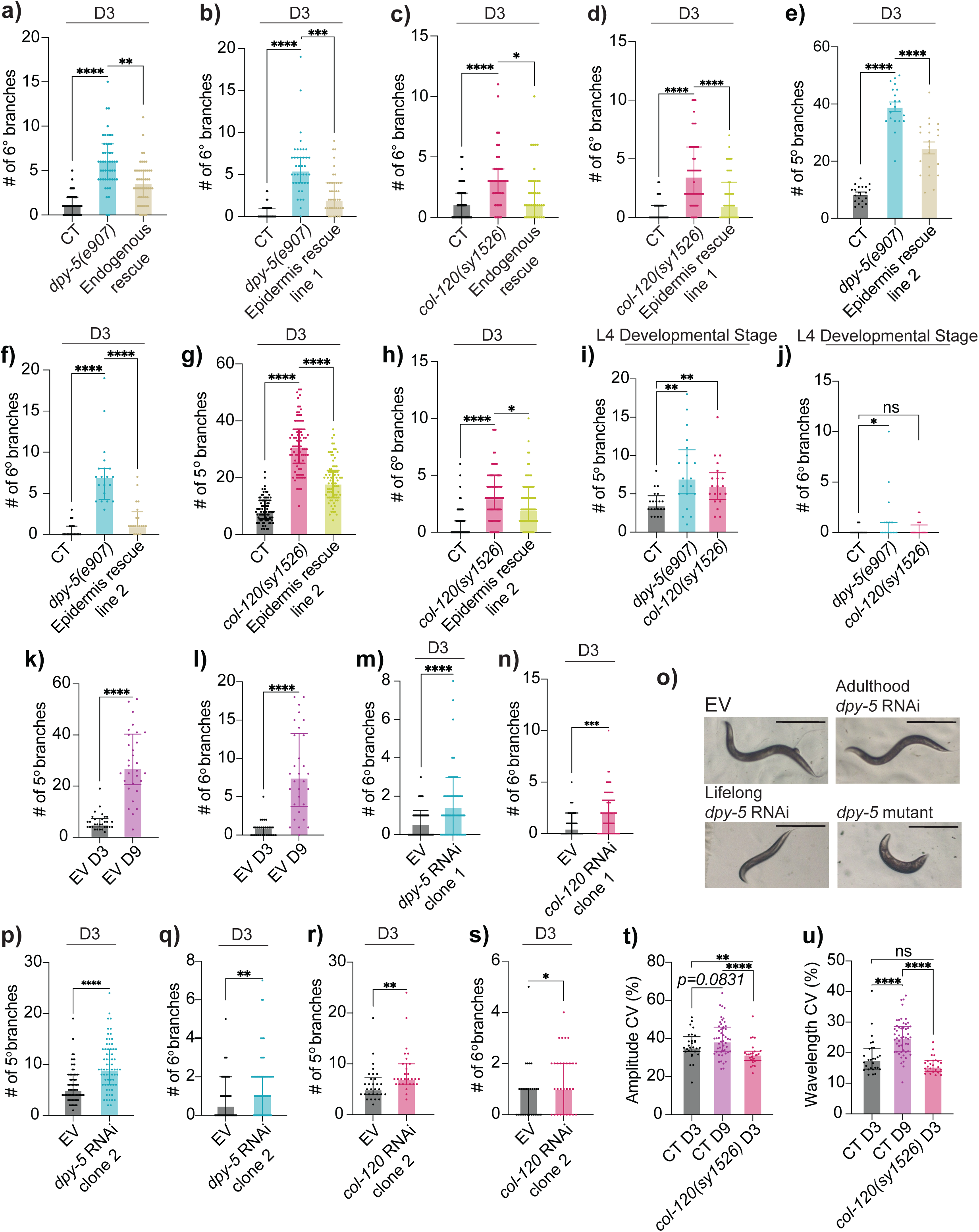
Characterization of loss of *dpy-5* and *col-120*. **(a)** Quantification of 6° dendrites in CT(n = 50), *dpy-5(e907)* mutant (n = 50), and endogenous promoter-driven rescue (n = 49). **(b)** 6° dendrites in CT, *dpy-5* mutant, and epidermis-specific rescue driven by *dpy-7p* (n = 60; line 1). **(c)** 6° dendrites in CT (n = 40), *col-120(sy1526)* (n = 38), and endogenous promoter-driven rescue (n = 38). **(d)** 6° dendrites in CT (n = 82), *col-120* mutant (n = 100), and epidermis-specific rescue driven by driven by *dpy-7p* and *col-19p* (n = 55; line 1). **(e-f)** 5° *(e)* and 6° *(f)* dendrites in CT, *dpy-5* mutant, and epidermis-specific rescue driven by *dpy-7p* (*n* = 20; line 2). **(g-h)** 5° *(g)* and 6° *(h)* dendrites in CT, *col-120* mutant, and epidermis-specific rescue driven by driven by *dpy-7p* and *col-19p* (n = 81; line 2). **(i-j)** 5° *(I)* and 6° *(j)* dendrites in CT, *dpy-5* mutant, and *col-120* mutant (n = 20 for all groups). **(k-l)** 5° *(k)* and 6° *(l)* dendrites in empty vector (EV) treated animals (n = 30 for both groups). **(m)** 5° dendrites in EV and *dpy-5* RNAi clone 1 treated animals (n = 88 for both groups). **(n)** 5° dendrites in EV and *col-120* RNAi clone 1 treated animals (n = 70 for both groups). **(o)** Representative images of adulthood EV-treated, adulthood *dpy-5* clone 1 RNAi-treated, lifelong *dpy-5* clone 1 RNAi-treated, and *dpy-5* mutant animals at D3. Scale bar = 500 μM. **(p-q)** 5° *(p)* and 6° *(q)* dendrites in EV and *dpy-5* RNAi clone 2 animals (n = 70 for both groups). **(r-s)** 5° *(r)* and 6° *(s)* dendrites in EV and *col-120* RNAi clone 2 animals (n = 30 for both groups). **(t-u)** Amplitude CV *(t)* and wavelength CV *(u)* in CT D3 (n = 28), CT D9 (n = 51), and *col-120* mutant (n = 25) at D3. ns- not significant* *p*<0.05, ** *p*<0.005, *** *p*<0.001, **** *p*<0.0001.

**Figure S6:**
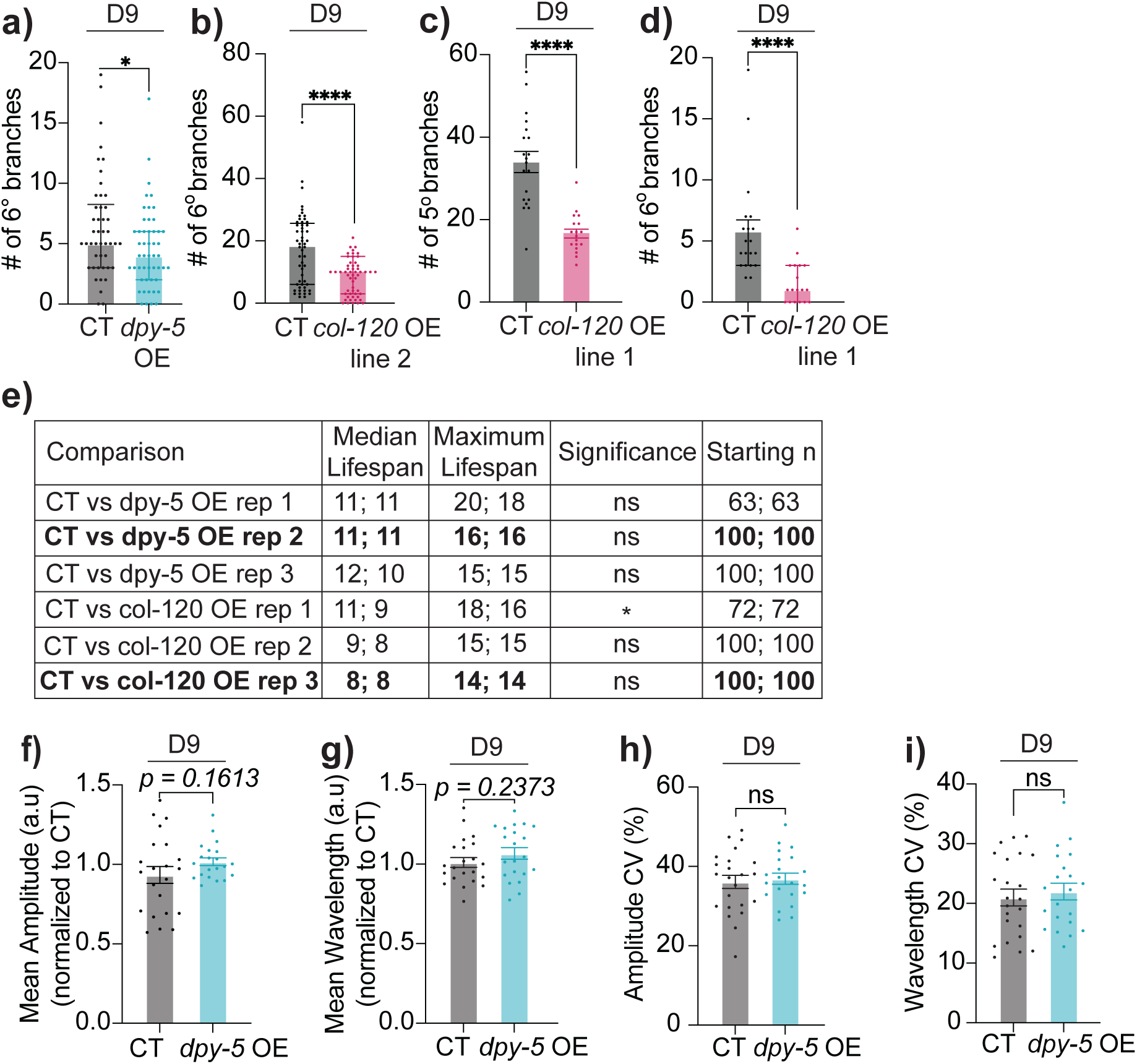
Overexpression of epidermal collagens. **(a)** Quantification of 6° dendrites in control (CT, n = 46) and *dpy-5* epidermis-specific overexpression driven by *col-19p* (n = 50). **(b)** 6° dendrites in CT (n = 53) and *col-120* epidermis-specific overexpression driven by *col-19p* (n =54; line 2). **(c-d)** 5° *(c)* and 6° *(d)* dendrites in CT (n = 20) and *col-120* epidermis-specific overexpression driven by *col-19p* (n = 18; line 1). **(e)** Summary of lifespan analysis of CT, *dpy-5* epidermis-specific overexpression, and *col-120* epidermis-specific overexpression line 1 animals without FUDR. Replicates in main figure indicated in bold. Same transgenic strains as in *(a)* and *(c-d)*. **(f-i)** Mean amplitude *(f)*, and wavelength *(g)* normalized to control, amplitude CV *(h)*, and wavelength CV *(i)* in CT (n = 23) in *dpy-5* overexpression (same transgenic strain as in *(a)*)(n = 21). ns- not significant, * *p*<0.05, *** *p*<0.001, **** *p*<0.0001.

**Figure S7:**
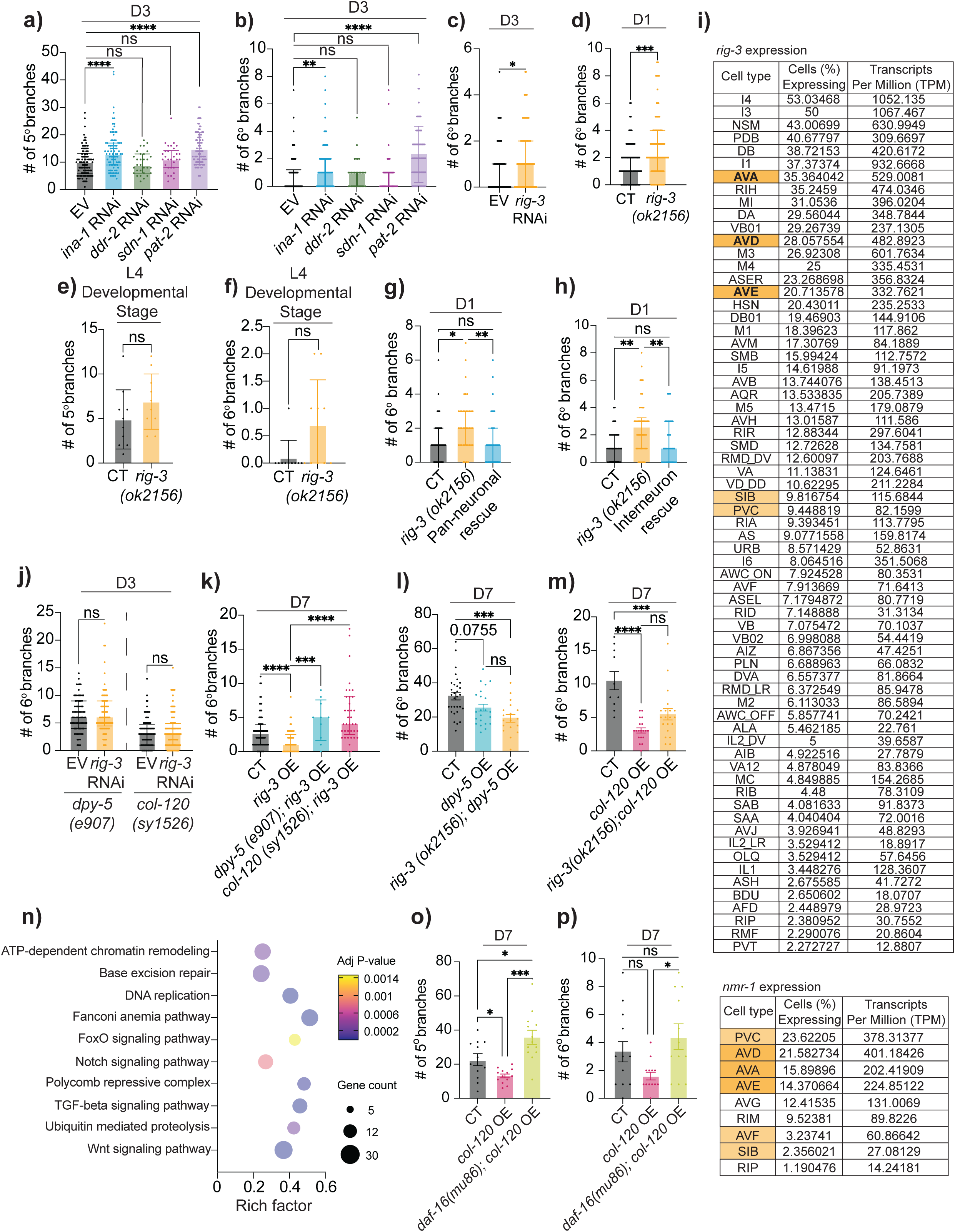
Interactions between *rig-3*/IgSF, collagen genes, and *daf-16*/FOXO. **(a-b)** Quantification of 5° *(a)* and 6° *(b)* dendrites in empty vector (EV, n = 208), *ina-1* RNAi (n = 80), *ddr-2* RNAi (n = 30), *sdn-1* RNAi (n = 30), *pat-2* RNAi treatment (n = 92). **(c)** 6° dendrites in EV and *rig-3* RNAi treated CT animals (*F49H12.4::GFP(wdIs51)*) *(n* = 84). **(d)** 6° dendrites in CT (*ser-2(3)p::GFP(lxyEx77)*) (n = 91) and *rig-3(ok2156)* mutant (n = 90). **(e-f)** 5° *(e)* and 6° *(f)* dendrites in CT and *rig-3* mutant (n = 10). (g) 6° dendrites in CT (n = 37), *rig-3* mutant (n = 40), and pan-neuronal rescue (n = 37) (driven by *unc-33p*). (h) 6° dendrites in CT (n = 41), *rig-3* mutant (n = 46), and interneuron rescue (n = 37) (driven by *nmr- 1p*). (i) Expression levels of *rig-3* and *nmr-1* in neuronal tissues of wild-type animals (Wormbase). (j) 6° dendrites in *dpy-5(e907)* mutant animals treated with EV (n = 84) or *rig-3* RNAi (n = 85), as well as in *col-120(sy1526)* mutant animals treated with EV (n = 88) or *rig-3* RNAi (n = 86). (k) 6° dendrites in CT (n = 76), *rig-3* pan-neuronal overexpression driven by *unc-33p* (n = 57), *rig-3* overexpression in *dpy-5* mutant (n = 10), and *rig-3* overexpression in *col-120* mutant (n = 37). (l) 6° dendrites in CT (n = 30), *dpy-5* epidermis-specific overexpression driven by *col-19p* (n = 22), and *dpy-5* overexpression in *rig-3* mutant (n = 18). (m) 6° dendrites in CT (n = 10), *col-120* epidermis-specific overexpression driven by *col-19p* (n = 19; line 1), and *col-120* overexpression in *rig-3* mutant (n = 20). (n) KEGG pathway analysis between control and *dpy-10* mutant animals (GSE19310). **(o-p)** 5° *(o)* and 6° *(p)* dendrites in CT (n = 12), *col-120* epidermis-specific overexpression driven by *col-19p* (n = 13; line 1), and *col-120* overexpression line 1 in *daf-16(mu86)* mutant background (n = 13). Experiment performed without FUDR. ns-not significant, * *p*<0.05, ** p<0.01, *** *p*<0.001, **** *p*<0.0001.

**Table S1:** Comparison of branching in PVDL and PVDR, and Dorsal, Ventral, Anterior, and Posterior sections of PVD neuron. Difference between medians for CT animals at D3 and D9 with FUDR, and D3 and D7 without FUDR. ns-not significant, * *p*<0.05, ** p<0.01, *** *p*<0.001, **** *p*<0.0001.

| Comparison | with FUDR |  |  | no FUDR |  |  |
| --- | --- | --- | --- | --- | --- | --- |
|  | CT D3 | CT D9 | Significance | CT D3 | CT D7 | Significance |
| Number of Right 5° branches | 2.0<br>(n = 58) | 20.0<br>(n = 58) | **** | 13.0<br>(n = 44) | 28.0<br>(n = 29) | **** |
| Number of Left 5° branches | 2.0<br>(n = 47) | 16.0<br>(n = 42) | **** | 10.50<br>(n = 50) | 27.00<br>(n = 41) | **** |
| Number of Right 6° branches | 1.0<br>(n = 58) | 5.5<br>(n = 58) | **** | 1.0<br>(n = 44) | 5.0<br>(n = 29) | **** |
| Number of Left 6° branches | 1.0<br>(n = 47) | 5.5<br>(n = 42) | **** | 1.0<br>(n = 50) | 5.0<br>(n = 41) | **** |
| Number of Dorsal menorahs | 16.5<br>(n = 10) | 20.0<br>(n = 11) | ns | 19.0<br>(n = 21) | 16.0<br>(n = 19) | ** |
| Number of Ventral menorahs | 17.0<br>(n = 20) | 16.0<br>(n = 14) | ns | 16.5<br>(n = 22) | 16.0<br>(n = 39) | ns |
| Number of Anterior menorahs | 12.5<br>(n = 30) | 13.0<br>(n = 25) | ns | 13.0<br>(n = 43) | 11.0<br>(n = 58) | *** |
| Number of Posterior menorahs | 5.0<br>(n = 30) | 5.0<br>(n = 25) | ns | 5.0<br>(n = 43) | 5.0<br>(n = 58) | ns |
| Number of Dorsal 4° branches | 66.0<br>(n = 10) | 78.0<br>(n = 11) | ns | 67.0<br>(n = 21) | 68.0<br>(n = 19) | ns |
| Number of Ventral 4° branches | 66.0<br>(n = 20) | 72.5<br>(n = 14) | ns | 77.0<br>(n = 22) | 75.0<br>(n = 39) | ns |
| Number of Anterior 4° branches | 40.0<br>(n = 30) | 44.0<br>(n = 25) | * | 47.0<br>(n = 43) | 45.0<br>(n = 58) | ns |
| Number of Posterior 4° branches | 26.0<br>(n = 30) | 29.0<br>(n = 25) | ns | 25.0<br>(n = 43) | 25.0<br>(n = 58) | ns |
| Number of Dorsal 5° branches/<br>Dorsal menorah | 0.5045<br>(n = 10) | 2.000<br>(n = 11) | *** | 0.0625<br>(n = 21) | 0.3158<br>(n = 19) | **** |
| Number of Ventral 5° branches/<br>Ventral menorah | 0.1563<br>(n = 20) | 0.3643<br>(n = 14) | **** | 0.0816<br>(n = 22) | 0.3571<br>(n = 39) | **** |
| Number of Anterior 5° branches/<br>Anterior menorah | 0.2020<br>(n = 30) | 1.1670<br>(n = 25) | **** | 0.0000<br>(n = 43) | 0.2000<br>(n = 58) | **** |
| Number of Posterior 5° branches/<br>Posterior menorah | 1.3750<br>(n = 30) | 3.0000<br>(n = 25) | **** | 0.1667<br>(n = 43) | 0.6333<br>(n = 58) | **** |
| Number of Dorsal 5° branches/<br>Dorsal 4° branches | 0.1257<br>(n = 10) | 0.4103<br>(n = 11) | *** | 0.0161<br>(n = 21) | 0.0641<br>(n = 19) | *** |
| Number of Ventral 5° branches/<br>Ventral 4° branches | 0.1563<br>(n = 20) | 0.3643<br>(n = 14) | **** | 0.0160<br>(n = 22) | 0.0806<br>(n = 39) | **** |
| Number of Anterior 5° branches/<br>Anterior 4° branches | 0.0519<br>(n = 30) | 0.3056<br>(n = 25) | **** | 0.0000<br>(n = 43) | 0.0454<br>(n = 58) | **** |
| Number of Posterior 5° branches/<br>Posterior 4° branches | 1.3750<br>(n = 30) | 3.0000<br>(n = 25) | **** | 0.0357<br>(n = 43) | 0.1200<br>(n = 58) | **** |
| Number of Dorsal 6° branches/<br>Dorsal menorah | 0.0000<br>(n = 10) | 0.2500<br>(n = 11) | ** | Refer to Figure 1 |  |  |
| Number of Ventral 6° branches/<br>Ventral menorah | 0.0000<br>(n = 20) | 0.1333<br>(n = 14) | *** |  |  |  |
| Number of Anterior 6° branches/<br>Anterior menorah | 0.0000<br>(n = 30) | 0.0909<br>(n = 25) | **** |  |  |  |
| Number of Posterior 6° branches/<br>Posterior menorah | 0.0000<br>(n = 30) | 0.4000<br>(n = 25) | **** |  |  |  |
| Number of Dorsal 6° branches/<br>Dorsal 4° branches | 0.0000<br>(n = 10) | 0.0635<br>(n = 11) | ** |  |  |  |
| Number of Ventral 6° branches/<br>Ventral 4° branches | 0.0000<br>(n = 20) | 0.0329<br>(n = 14) | ** |  |  |  |
| Number of Anterior 6° branches/<br>Anterior 4° branches | 0.0000<br>(n = 30) | 0.0238<br>(n = 25) | **** |  |  |  |
| Number of Posterior 6° branches/<br>Posterior 4° branches | 0.0000<br>(n = 30) | 0.0689<br>(n = 25) | **** |  |  |  |

**Table S2:**
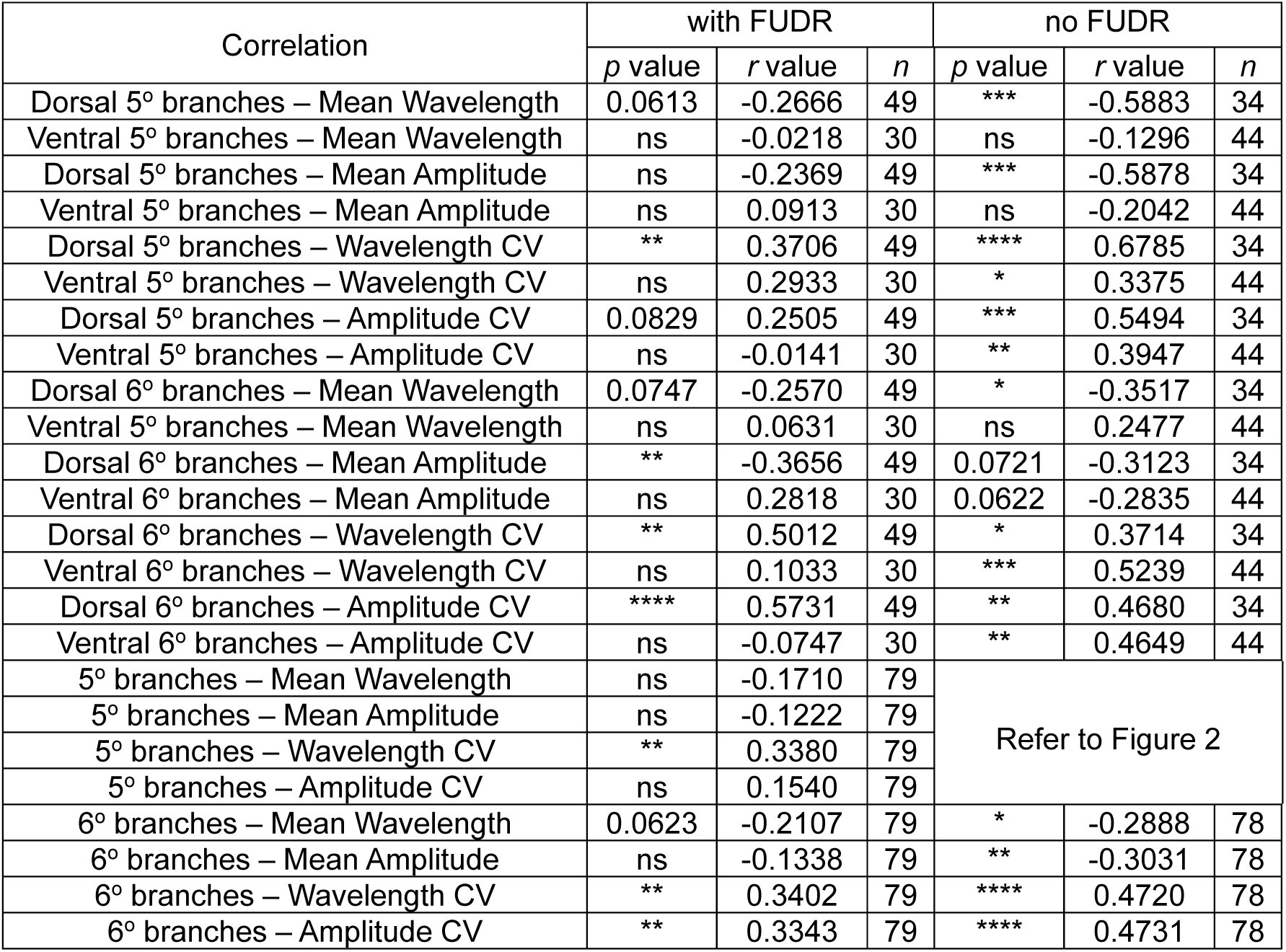
Correlation between excessive PVD dendritic branching and proprioceptive deficits. D3 and D9 CT animals were assayed and combined for analysis when FUDR was used for age synchronization; Data from D3 and D7 CT animals were collected when assays were conducted without FUDR. ns-not significant,* *p*<0.05, ** p<0.01, *** *p*<0.001, **** *p*<0.0001.

## Notes

### Competing Interest Statement

The authors have declared no competing interest.

### Summary of Updates

We have revised the title of our manuscript

